# Intranasal Delivery of Hypoxia-Primed Wharton’s Jelly MSC-Derived sEVs Reprograms Neuroimmune Signalling and Ameliorates Behavioural Deficits in a Valproic Acid–Induced Mouse Model of Autism Spectrum Disorder

**DOI:** 10.64898/2026.05.24.721255

**Authors:** Shruti Mahapatra, Bhanu Teja, Ritesh Kumar Netam, Neena Malhotra, Tulika Seth, Sheffali Gulati, Sujata Mohanty

## Abstract

**Background:** Beyond disrupted neurodevelopment, autism spectrum disorder (ASD) encompasses progressive neurodegenerative alterations that remain refractory to standard of care behavioral therapies, highlighting the need for biologically targeted adjunctive interventions. Although, Mesenchymal stem cell (MSC) derived small extracellular vesicles (sEVs) have emerged as potent neuroprotective agents; however, the optimal MSC tissue source and strategies enhancing their efficacy for clinical translation remain undefined. Hypoxic preconditioning potentiates their therapeutic effects, however its specific impact on ASD-related impairments is yet to be established. This study conducts a head to head evaluation of sEVs from bone marrow (BM) and Whartons jelly (WJ) MSCs under normoxic and hypoxic conditions to delineate their comparative neuroregenerative efficacy in ASD.

**Methods:** We investigated the neuroprotective efficacy of sEVs derived from human tissue-specific mesenchymal stem cells (BM/WJ) cultured under normoxic (21% O₂) and hypoxic (1% O₂) conditions. *In vitro* assays evaluated sEV uptake, cell proliferation, neurodifferentiation, mitochondrial health, and immunomodulatory effects using neural stem cell (C17.2) and microglial (N9) cell lines. For *in vivo* studies, ASD-like phenotypes were induced in C57BL/6 mice via prenatal exposure to valproic acid (600 mg/kg), followed by the biodistribution of fluorescently labeled hypoxia-primed WJ MSCs sEVs (WJ-H-sEVs) following intranasal administration was assessed using IVIS imaging. Behavioural outcomes after intranasal treatment were evaluated. Additionally, post-mortem brain tissues were analyzed for neuroinflammation, oxidative stress, synaptic integrity, and downstream signalling pathways using immunofluorescence, Western blotting, and qRT-PCR, while systemic cytokine pro-inflammatory and anti-inflammatory levels were quantified by ELISA. Mechanistic involvement of Nrf2 and NF-κB signaling pathways was examined using pharmacological inhibitors.

**Results:** Comparative analyses revealed that hypoxic priming significantly enhanced sEV yield and enriched neuroprotective miRNAs, with WJ-H-sEVs exhibiting elevated levels of miR-125b-5p, miR-21a-5p, and miR-145a-5p. Consistent with this enrichment, WJ-H-sEVs demonstrated superior cellular uptake as observed in neural stem cells and microglia, enhanced neurodifferentiation, robust antioxidant activity, effective immunomodulation, and improved mitochondrial health, thereby outperforming their normoxic and BM-derived counterparts. Intranasally administered WJ-H-sEVs efficiently localized to the brain within 12 h and significantly improved anxiety-like behavior, recognition memory, and spatial learning in ASD mice. These effects were accompanied by reduced neuroinflammation and oxidative damage, enhanced neuronal survival, decreased serum IL-6 levels, and increased IL-10 and TGF-β. Mechanistically, therapeutic benefits were mediated through activation of the Nrf2 antioxidant pathway and suppression of NF-κB and JAK-STAT signaling.

**Conclusion:** In this study, hypoxia emerges as a clinically exploitable priming strategy to enhance the neuroregenerative performance of tissue-specific MSC-derived sEVs, with potential applicability across MSC sources and other neurological disorders. Moreover, intranasal delivery of WJ-H-sEVs demonstrates translational feasibility, reinforcing their potential as an adjunctive, cell-free, pediatric-friendly neurotherapeutic modality for ASD management.

**Graphical Abstract:** 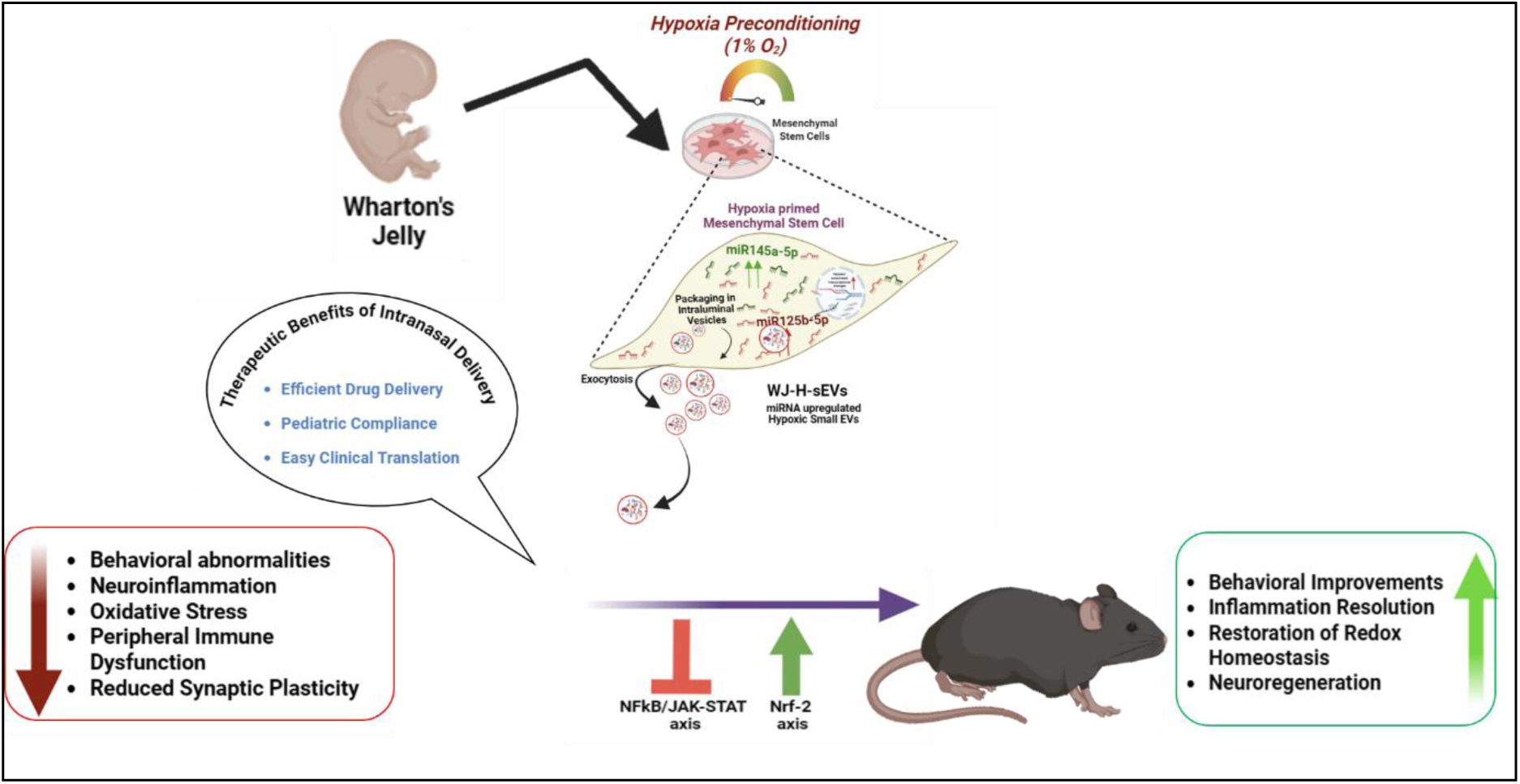

## Introduction

ASD is neurodevelopmental condition characterized by early impairments in social communication, restricted interests, and repetitive sensory-motor behaviors^1,2^. Symptoms typically emerge within the first two to three years of life, and ASD can be reliably diagnosed as early as 18–24 months of age, although many individuals are diagnosed later in childhood depending on symptom severity. The diagnosis primarily follows DSM-5 (Diagnostic and Statistical Manual of Mental Disorders, Fifth Edition)^3^ and ICD-11 (International Classification of Diseases, 11th Revision) criteria^4^, and the clinical presentation is highly heterogenous^5^, ranging from subtle social difficulties to prominent repetitive behaviour and speech patterns, marked deficits in verbal and non-verbal communication^6^, and co-occurring anxiety^7^, ADHD (attention-deficit/hyperactivity disorder)^8^, sleep disturbances^9^, intellectual disability^10^, and gastrointestinal problems^11^. Although ASD shows high heritability, identified genetic variants account for only a fraction of overall risk and explain less than 10% of cases linked to single-gene syndromes such as tuberous sclerosis, fragile X, or Rett syndrome, indicating that most ASD arises from the combined effects of multiple genetic variants and environmental influences that trigger pathological physiological changes in genetically susceptible individuals^12–14^.

A 2021 *Lancet* Commission report highlighted ASD is a growing public health concern, affecting an estimated 78 million people, with recent 2023 CDC data, indicating a prevalence of 1 in 36 children in the USA^15^. In India, a recent comprehensive study indicated that approximately 1 in 89 children have ASD^16^, highlighting ASD as a growing public health impact and the need for disease-modifying therapies. Despite the escalating prevalence and substantial the financial and social impact of this condition^17,18^, clinical management remains largely supportive and symptomatic, relying on behavioural interventions^19^, speech and occupational therapy^20^, and limited pharmacological agents^21^ targeting associated features such as irritability or anxiety. While behavioral interventions can be initiated as early as 24 months, pharmacological therapies for ASD-associated symptoms are generally approved only for children older than five years, despite compelling evidence that interventions introduced during earlier neurodevelopmental windows may confer more robust and durable benefits. This therapeutic gap highlights the critical need for interventions that can be administered during early neurodevelopmental stages and directly address dysregulated neurobiological mechanisms, modify disease trajectories, and improve long-term functional outcomes.

While ASD is classically defined as a neurodevelopmental disorder, accumulating longitudinal, neuropathological, and molecular data indicate that, it also acquires neurodegenerative-like features that evolve over time^22–24^. Rett syndrome is a genetically defined neurodevelopmental disorder that predominantly affects girls and shares core clinical and biological features with ASD, caused primarily by mutations in *MECP2*^25^, provides a well-established precedent in which a developmental origin is accompanied by progressive neuronal dysfunction driven by oxidative stress and sustained low-grade inflammation are mechanistically intertwined rather than acting as independent processes, a process known as “OxInflammation”^26–29^.

Substantial evidence now unequivocally points to analogous oxinflammatory signatures in ASD, including neuroinflammation, immune activation, oxidative stress and neuronal dysfunction. Post-mortem investigations show chronic microgliosis and astrogliosis^30,31^, while neuroimaging reveals dysfunction of cortico-limbic and fronto-striatal circuits^32^. Multiple studies report decreased enzymatic antioxidants^33^ and anti-inflammatory cytokines^34^, together with increased DNA, lipid, and protein oxidation products^35^ and elevated pro-inflammatory cytokines^36^ in brain and peripheral fluids. These oxidative stress and inflammatory markers in peripheral biofluids correlate with ASD severity, including social-communication deficits and behavioural disturbances. These findings indicate that oxinflammation functionally contributes to core ASD symptoms, mirroring neurodegenerative diseases like Parkinson’s, Alzheimer’s, and amyotrophic lateral sclerosis (ALS), where chronic immune/redox dysregulation drives clinical progression. This suggests early neurodevelopmental perturbations in ASD are compounded by persistent oxinflammatory cascades, yielding a neurodegenerative phenotype, though the precise sequence of these events remains incompletely understood. Given this complex pathophysiology and limitations of current treatments, biologic approaches targeting cellular and molecular underpinnings in ASD represent a rational strategy to address this major therapeutic gap.

Over the past two decades, regenerative medicine has shifted from mesenchymal stem cells (MSCs) to MSC-derived small extracellular vesicles (sEVs) as a compelling cell-free modality that targets the underlying complex pathophysiology to reshape the dysregulated molecular landscape of neurological disorders including neurodegenerative conditions (Parkinson’s, Alzheimer’s, ALS) and neurodevelopmental disorders (cerebral palsy, Rett syndrome, hypoxic-ischemic encephalopathy, ASD, perinatal brain injury). MSC-sEVs can traverse biological barriers and deliver regulatory cargoes, particularly miRNAs, along with proteins and mRNAs, that modulate recipient cell signaling and functional states. Through coordinated regulation of pathways involved in neuroinflammation, oxidative stress, and cellular survival such as NF-κB, JAK/STAT, PI3K/AKT, MAPK, and Nrf2/HO-1, MSC-sEVs have been shown to exert context-dependent immunomodulatory and neuroprotective effects in both neurodegenerative and neurodevelopmental disorders, including ASD.

Nevertheless, several critical questions remain unresolved before MSC-derived small extracellular vesicles (MSC-sEVs) can be rationally advanced as an adjunctive therapy for ASD. First, the ideal MSC tissue source for generating maximally neuroprotective sEVs is undefined, with adult (BM) and perinatal (WJ) MSCs exhibiting distinct immunophenotypes and secretomes that may influence therapeutic consistency. Second, both sEV yield and neuroprotective potency are strongly shaped by the biogenesis microenvironment, particularly oxygen tension. While hypoxic preconditioning enhances sEV yield and regenerative miRNA enrichment, its ability to augment anti-inflammatory and neuroprotective effects in ASD has not been systematically assessed. Third, the endogenous miRNA cargo mediating sEV-driven modulation of ASD pathophysiology remains poorly characterized. Fourth, the optimal route of administration for delivering MSC-sEVs in pediatric ASD populations remains inadequately defined.

Against this backdrop, the present study thoroughly compares the neuroprotective efficacy of sEVs derived from BM and WJ-MSCs under normoxic (21% O₂) conditions to identify the most effective MSC source. Subsequently, hypoxic preconditioning (1% O₂) is evaluated as a strategy to further bolster the neuroprotective potential of tissue specific sEVs. These effects are validated through comprehensive *in vitro* and *in vivo* functional assessments, alongside mechanistic interrogation of the neuroimmune and redox signaling pathways underlying sEV-mediated therapeutic effects. Collectively, this study identifies the most suitable MSC source for clinically translatable sEV production and establishes hypoxic priming as a viable strategy to potentiate the neuroprotective efficacy of MSC-sEV–based, cell-free therapies for ASD. Furthermore, intranasal delivery represents an efficient and clinically practical approach for targeting EVs to brain tissue.

## Results

### Low oxygen culture conditions elevate sEV production and neuroprotective miRNA levels in tissue specific MSCs

In this study we used human derived BM-MSCs and WJ-MSCs fulfilled ISCT^37^ criteria by displaying spindle-shaped, fibroblast-like morphology under both normoxic and hypoxic culture, robust plastic adherence, and tri-lineage differentiation into osteogenic, adipogenic, and chondrogenic lineages as evidenced by Alizarin Red, Oil Red O, and Alcian Blue staining, respectively (**Fig.S1a–b**). Flow cytometry confirmed high expression of canonical MSC markers CD105, CD29, CD90, CD73, and HLA-I, with minimal/absent CD34, CD45, and HLA-II, in both BM- and WJ-MSC preparations (**Fig. S1c**).

Following 24 h exposure to normoxia (21% O₂) or hypoxia (1% O₂), sEVs were isolated from the collected conditioned media of tissue specific MSCs using sucrose density based ultracentrifugation. NTA showed that with modal diameters of 101.9 nm (mean 101.9 ±14.1nm) BM-MSC-sEVs normoxia and 88.0 nm for BM- (mean 80 ±14.1nm) hypoxia and WJ-MSC-sEVs under normoxia, 123.5 nm (mean 123.5 ± 6.23 nm) and 93.2 nm (mean 93.2 ±14.1nm) under hypoxia, respectively (**Fig. 1a**).

**Fig. 1.**
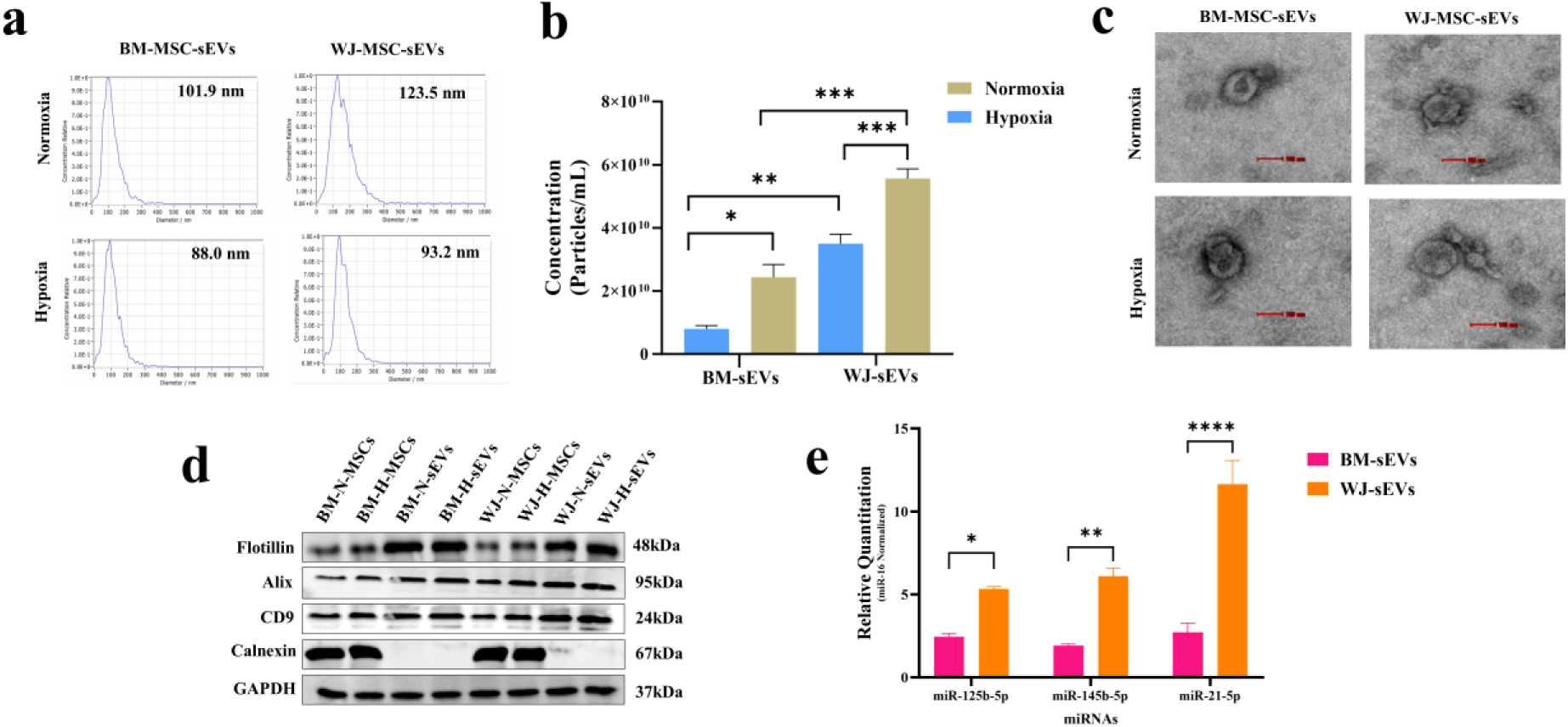
Characterization of human tissue-specific MSCs (BM/WJ) secreted sEVs under normoxic (21% O₂) and hypoxic (1% O₂) conditions, and comparison of neuroprotective miRNA expression profiles. In accordance with MISEV2023 recommendations, multiple complementary approaches were used to confirm the identity, purity, and functional relevance of the isolated sEVs. (a) Particle size distribution assessed by NTA. (b) Quantification of sEV concentrations derived from BM- and WJ-MSCs cultured under normoxic and hypoxic conditions. (c) Quantitative PCR analysis of miR-125b-5p, miR-145b-5p, and miR-21-5p in normoxic and hypoxic sEVs from BM- and WJ-MSCs. (d) Transmission electron microscopy images showing typical cup-shaped vesicles (Scale bar: 100 nm). (e) Immunoblot analysis confirming the presence of canonical sEV markers (Flotillin, CD81, TSG101) and the housekeeping control GAPDH and the absence of the negative marker calnexin. (f) Data are presented as mean ± SD (n = 3); *p < 0.05, **p < 0.01, ***p < 0.001, ****p < 0.0001

Hypoxic priming markedly enhanced sEV production from both MSC sources, with particle concentrations increasing from 2 × 10¹⁰ ± 0.62 × 10¹⁰ to 4.5 × 10¹⁰ ± 0.20 × 10¹⁰ particles/mL for BM-MSCs (2.3-fold, p < 0.05) and from 4 × 10¹⁰ ± 0.18 × 10¹⁰ to 7.5 × 10¹⁰ ± 0.32 × 10¹⁰ particles/mL for WJ-MSCs (1.9-fold, p < 0.001), highlighting WJ-MSCs as a comparatively higher-yield source of therapeutic sEVs (**Fig. 1b**), highlighting WJ-MSCs as a high-yield source for therapeutic sEV production. TEM confirmed abundant cup-shaped vesicles of 30–150 nm for all groups (**Fig. 1c**), while Western blotting showed enrichment of canonical sEV markers Flotillin, Alix, and CD9 in sEV fractions and absence of the ER protein Calnexin, consistent with Minimal information for studies of extracellular vesicles (MISEV 2023)^38^ criteria for small EV preparations (**Fig. 1d**).

Previous work has shown that the predominant functional RNA cargo in sEVs consists of miRNAs, which can reprogram cellular phenotypes. Others and our group have further demonstrated that physiological cues applied to parent cells, including hypoxia, influence the miRNA packaging profile of sEVs^39–41^. Hence, next we quantified the expression of three miRNAs with known neuroprotective and immunomodulatory roles, miR-125b-5p, miR-145-5p, and miR-21-5p, in hypoxia-primed sEVs. qRT-PCR revealed significantly higher levels of all three miRNAs in WJ-sEVs than in BM-sEVs under hypoxia, with approximately 3.8-fold enrichment of miR-125b-5p (p < 0.05), 3 fold enrichment of miR-145-5p (p < 0.01), and 9.2-fold enrichment of miR-21-5p (p < 0.0001) (**Fig. 1e**).

Overall, MSC-sEV preparations contained high levels of miR-21-5p, consistent with prior reports that this miRNA constitutes a major fraction of MSC-exosomal miRNA cargo^42^. For instance, miR-21-5p has been shown to promote neuronal survival and limits apoptosis after CNS injury, largely via repression of PTEN and activation of the PI3K–AKT pathway^43^, as well as modulation of PDCD4 and FasL–caspase signalling pathway^44^, thereby facilitating functional recovery. Likewise, miR-125 is significantly enriched following hypoxic preconditioning and have been shown to dampen microglial activation and neuroinflammation and to support neurovascular regeneration, in part by targeting pro-inflammatory mediators such as IRF5^45^ and components of the NF-κB pathway, and was also reported to enhance acidosis resilience after cerebral ischaemia–reperfusion by suppressing ASIC1 protein expression^46^. In addition, miR-145 delivered via MSC-derived exosomes exerts anti-inflammatory, anti-apoptotic and anti-fibrotic effects in part through direct targeting of FOXO1 in cerebral ischaemia reperfusion injury^47^ and functional recovery after spinal cord injury via regulation of Plexin-A2 signalling^48^. It has also been linked to regulation of TLR-4& NLRP3^49^ inflammasome activity, thereby modulating NF-κB^49–51^ signalling leading to promotion of M2 microglia^51^ and modulate endothelial cell function by targeting ROCK1^52^. Consistent with these reports, we observed pronounced enrichment of miR-21-5p, miR-125b-5p, and miR-145-5p, with the most robust enrichment detected in WJ-sEVs under both normoxic and hypoxic conditions in the present study.

Taken together, these findings indicate that WJ-MSCs preserve core MSC characteristics under low oxygen while producing substantially higher quantities of sEVs and packaging more neuroprotective miRNAs than BM-MSCs. This is consistent with previous reports showing that WJ-MSCs are superior EV producers and supports their use as a favourable tissue source for translational applications^53^, while hypoxia acts as a broadly applicable priming strategy that boosts sEV biogenesis and enriches functionally relevant miRNA cargo via HIF-dependent modulation of endosomal trafficking and RNA sorting^54–60^.

### WJ-H-sEVs demonstrate superior neuroregenerative properties *in vitro*

To assess the differences in neuroregenerative potential arsing from tissue source and hypoxia priming,we evaluated sEV uptake, proliferation, migration, and differentiation in C17.2 mouse neural progenitor cells.

Effective vesicle internalisation is a prerequisite for downstream biological activity. Hypoxic preconditioning significantly enhanced sEV uptake compared with normoxic counterparts, irrespective of tissue source^61,62^. Notably, WJ-MSC-derived sEVs exhibited markedly greater internalisation than BM-MSC-sEVs, an effect that was further amplified under hypoxic conditions (**Fig. 2a,b**). Quantitative integrated density analysis confirmed that WJ-H-sEVs were internalised at significantly higher levels than BM-H-sEVs (p < 0.001), WJ-N-sEVs (p < 0.01), and BM-N-sEVs (p < 0.0001). Additionally, WJ-N-sEVs outperformed BM-N-sEVs (p < 0.01), while BM-H-sEVs showed greater uptake than BM-N-sEVs (p < 0.05) and untreated controls (p < 0.01). Overall, WJ-sEVs demonstrated superior internalisation compared with BM-sEVs (WJ-H > BM-H; WJ-N > BM-N), with hypoxia significantly enhancing uptake within each tissue source (WJ-H vs WJ-N, p < 0.01; BM-H vs BM-N, p < 0.05). These findings are consistent with reports that hypoxic conditioning enriches sEV surface tetraspanins (CD9, CD63, CD81), integrins, heat-shock proteins, phosphatidylserine exposure, and membrane sphingolipid and cholesterol content, thereby enhancing membrane fluidity, stability, and receptor-mediated endocytosis. Such hypoxia-induced surface and lipidomic remodeling likely underlies the preferential docking and uptake of WJ-H-sEVs by neural progenitors, although the precise receptor–ligand interactions warrant further investigation.

**Fig. 2.**
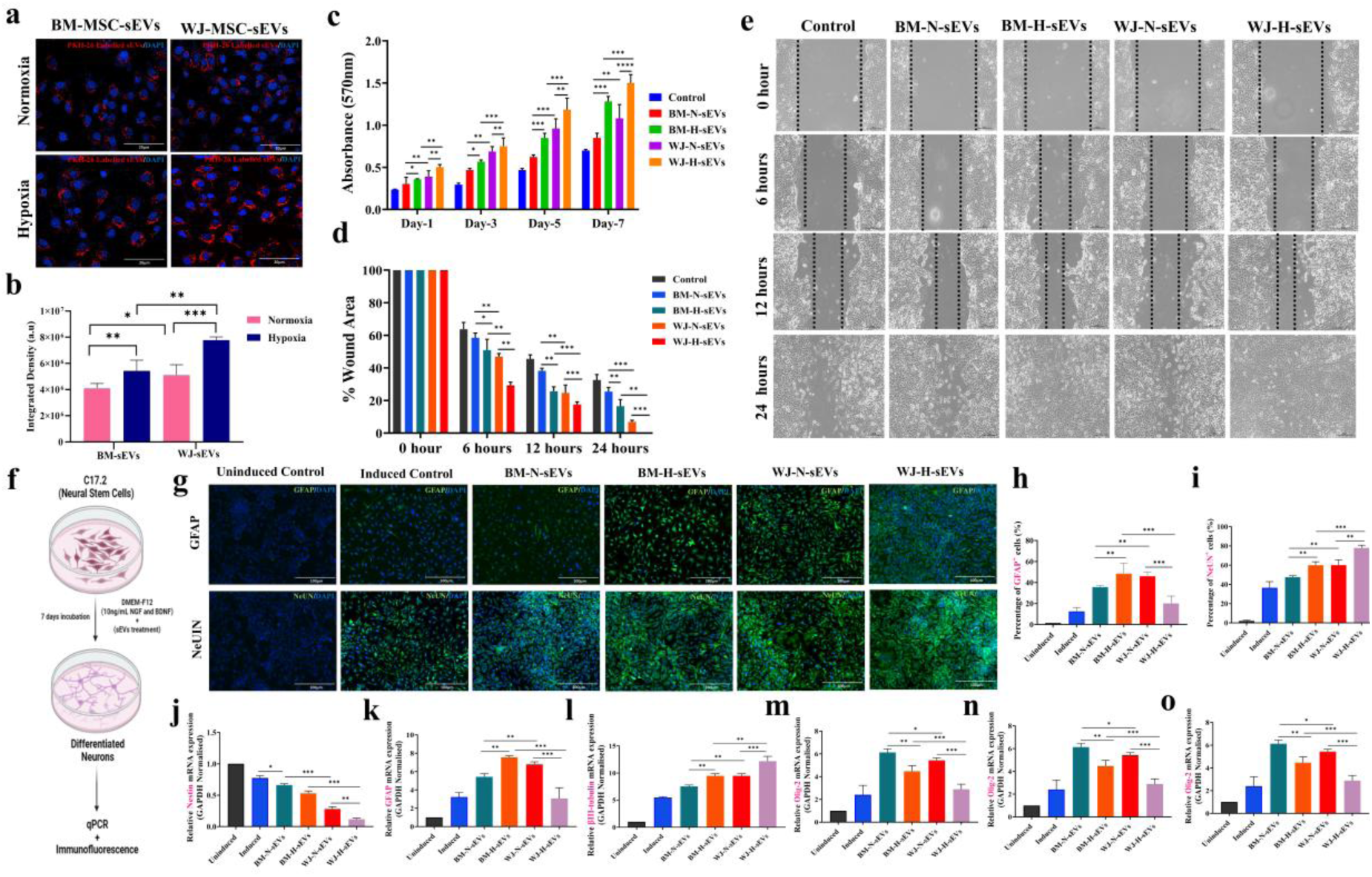
Hypoxia heightens WJ-sEVs internalization and neuroregenerative effects compared to normoxic counterpart and BM-sEVs in C17.2 neural stem cells. (a) PKH26-labeled tissue specific sEVs uptake by C17.2 neural stem cells under normoxia and hypoxia, showing increased internalization of WJ-H-sEVs as indicated by higher red fluorescence intensity with quantitative integrated density analysis (ii) (Scale bar: 20 µm). (B) Effect of normoxic and hypoxic BM- and WJ-MSC-sEVs on neural stem cell proliferation and migration; MTT assay absorbance at 570 nm on days 1, 3, 5, and 7 after sEV treatment is shown in (i), quantification of wound closure (%) at 0, 6, 12, and 24 h following scratch injury in the presence of the indicated sEVs is shown in (ii), and representative phase-contrast wound-healing images are shown in (iii) (Scale bar: 100 µm). (C) sEV-induced differentiation of C17.2 neural stem cells into neurons and astrocytes; the schematic of the differentiation protocol using DMEM/F12 containing 10 ng/mL NGF and BDNF and 30 µg sEVs protein for 7 days, followed by analysis of differentiation markers by immunofluorescence and qRT-PCR, is shown in (i). Representative immunofluorescence images for neuronal marker NeuN and astrocytic marker GFAP after treatment with normoxic or hypoxic BM- and WJ-MSC-sEVs are shown in (ii) (Scale bar: 100 µm), quantification of NeuN⁺ and GFAP⁺ cells (%) is shown in (iii) and (iv), respectively, and relative mRNA expression of Nestin, βIII-tubulin, Olig2, NeuN, MAP2 and GFAP by qPCR is shown in (v-x). Data are presented as mean ± SD (n = 3); and analyzed by one-way ANOVA with Bonferroni post hoc test; *p < 0.05, **p < 0.01, ***p < 0.001, ****p < 0.0001. Schematic illustration was created with BioRender.com.

Next, we examined the proliferative effects of sEVs on C17.2 cells over a 7-day period (**Fig. 2c**). By day 7, proliferation followed a clear hierarchy: BM-N-sEVs (p < 0.05) < BM-H-sEVs (p < 0.001) < WJ-N-sEVs (p < 0.0001) < WJ-H-sEVs (p < 0.0001). Hypoxic priming significantly enhanced the proliferative efficacy of BM-sEVs, with BM-H-sEVs inducing greater proliferation than BM-N-sEVs on day 5 (p < 0.01) and day 7 (p < 0.05). Importantly, WJ-derived sEVs consistently outperformed BM-derived sEVs at all time points. This source-dependent superiority was evident as early as day 1 (WJ-H-sEVs > BM-H-sEVs, p < 0.01) and remained significant on day 3 (WJ-N-sEVs > BM-N-sEVs, p < 0.05), day 5 (WJ-H-sEVs > BM-H-sEVs, p < 0.01; WJ-N-sEVs > BM-N-sEVs, p < 0.001), and day 7 (WJ-H-sEVs > BM-H-sEVs, p < 0.001; WJ-N-sEVs > BM-N-sEVs, p < 0.0001). These results indicate that WJ-MSC-sEVs, particularly under hypoxic priming, possess enhanced mitogenic capacity for neural progenitors, consistent with reports demonstrating enrichment of PI3K/AKT-, ERK/MAPK-, and Wnt/β-catenin-activating miRNAs and growth factors in hypoxia-conditioned WJ-MSC secretomes.

Scratch wound assay further demonstrated that sEV treatment significantly enhanced C17.2 migratory capacity compared with untreated controls (**Fig. 2d,e**). At 6 h, the WJ-H-sEV-treated group exhibited the greatest wound closure, with only 75 ± 3% of the wound area remaining compared with 90 ± 3% in controls. This effect was significantly greater than that observed with BM-H-sEVs (85 ± 3%, p < 0.001), WJ-N-sEVs (82 ± 2.5%, p < 0.01), and BM-N-sEVs (88 ± 2.5%, p < 0.0001). By 12 h, WJ-H-sEV treatment further reduced the residual wound area to 45 ± 4%, significantly outperforming rate/reduction BM-H-sEVs (60 ± 4%, p < 0.01), WJ-N-sEVs (55 ± 3.5%, p < 0.05), and BM-N-sEVs (70 ± 3.5%, p < 0.001). At 24 h, WJ-H-sEVs achieved near-complete closure (15 ± 3%), which was significantly superior to BM-H-sEVs (25 ± 3.5%, p < 0.01), WJ-N-sEVs (22 ± 3%, p < 0.05), and BM-N-sEVs (35 ± 4%, p < 0.0001). Across all time points, WJ-sEVs consistently outperformed BM-sEVs (WJ-H > BM-H; WJ-N > BM-N), while hypoxic priming significantly enhanced migratory efficacy within each tissue source. These findings align with studies demonstrating that hypoxia-conditioned WJ-MSC-sEVs promote cytoskeletal remodeling and chemotactic migration via CXCL12–CXCR4, PI3K/AKT, and ERK/MAPK signaling pathways.

qRT-PCR To evaluate lineage specification, C17.2 cells were cultured under differentiation conditions in the presence of sEVs for 7 days, demonstrating that WJ-H-sEVs significantly downregulated progenitor markers Nestin (p < 0.001 vs control; p < 0.01 vs BM-H-sEVs) and Olig2 (p < 0.01 vs control; p < 0.05 vs WJ-N-sEVs), while robustly upregulating neuronal markers βIII-tubulin (p < 0.0001 vs all groups), NeuN (p < 0.0001 vs BM-H-sEVs and normoxic groups), and MAP2 (p < 0.0001 vs control; p < 0.001 vs BM-N-sEVs) (**Fig. 2j–o**). GFAP mRNA expression remained significantly lower in WJ-H-sEV-treated cells than in BM-H-sEV-treated cells (p < 0.01), further confirming suppression of astroglial specification.

Immunofluoresence analysis corroborated these findings (Fig. 2f–o).Under these conditions, WJ-H-sEVs favored multiple neuronal lineage commitment while limiting astroglial expansion as evidenced by NeUN and GFAP (expression of these proteins). Immunofluorescence analysis revealed that WJ-H-sEV treatment yielded the highest proportion of NeuN⁺ neurons (p < 0.0001 vs BM-H-sEVs; p < 0.001 vs WJ-N-sEVs; p < 0.0001 vs BM-N-sEVs and control) and a concomitant reduction in GFAP⁺ astrocytes (p < 0.01 vs BM-H-sEVs; p < 0.05 vs WJ-N-sEVs; p < 0.001 vs control) (Fig. 2g–i).

Based on the above two experiments we found that multilineage neuronal expression his shift indicates preferential neuronal maturation over astroglial differentiation.

In contrast, BM-H-sEVs supported both neuronal and astroglial differentiation, consistent with reports that BM-MSC-sEVs preferentially activate gliogenic pathways, including STAT3, NF-κB, Notch, and SMAD signaling. Normoxic sEVs from either source supported mixed lineage outcomes. Thus, a clear source- and oxygenation-dependent divergence emerged, with BM-H-sEVs predominantly promoting gliogenesis, whereas WJ-H-sEVs strongly favored neuronal lineage commitment. This divergence mirrors emerging evidence that hypoxia-primed WJ-MSC-sEVs are enriched in neurogenic miRNAs (e.g., miR-21, miR-124, miR-132) and neurotrophic factors (e.g., BDNF, VEGF), which suppress Notch- and STAT3-dependent astrocytic differentiation while activating PI3K/AKT, ERK/MAPK, and Wnt/β-catenin signaling.

Overall, these data demonstrate that hypoxia-primed WJ-MSC-sEVs are internalised at an early time point by neural progenitors cells and exert superior pro-proliferative, pro-migratory, and neurogenic effects compared with normoxic and BM-derived sEVs. The observed source- and conditioning-dependent divergence underscores the importance of MSC origin and microenvironmental cues. This further provides a strong rationale for prioritising WJ-H-sEVs as a neuroregenerative modality for ASD and related neurodevelopmental disorders.

### Hypoxia enhances the immunoregulatory capacity of WJ-sEVs leading to anti-inflammatory reprogramming of macrophages and microglia

Sustained microglial activation and a bias toward the pro-inflammatory M1 phenotype have been repeatedly reported in ASD patient brains and relevant animal models, where elevated expression of iNOS, IL-1β, and TNF-α correlates with aberrant synaptic pruning and altered neurodevelopmental trajectories. Given the enhanced neuroregenerative potency of hypoxia-primed WJ-MSC-derived sEVs observed in neural progenitors, we next investigated whether these vesicles could more effectively modulate microglial polarization under inflammatory conditions.

To assess sEV internalization by microglia, PKH26-labeled sEV uptake was evaluated in N9 mouse microglial cells under normoxic and hypoxic conditions (**Fig. 3a,b**). Hypoxic priming significantly enhanced sEV uptake relative to normoxic counterparts, irrespective of tissue source. Notably, WJ-MSC-derived sEVs exhibited substantially higher internalization than BM-MSC-sEVs, an effect that was further amplified under hypoxia. Quantitative integrated density analysis confirmed that WJ-H-sEVs were internalized at significantly higher levels than BM-H-sEVs (p < 0.001), WJ-N-sEVs (p < 0.01), and BM-N-sEVs (p < 0.0001) fold change. These findings indicate that hypoxia conditioning and tissue origin cooperatively enhance sEV–microglia interactions, consistent with evidence that hypoxia can increase exosome uptake by target cells via altered integrin and heparan sulfate proteoglycan–dependent endocytosis.

**Figure 3.**
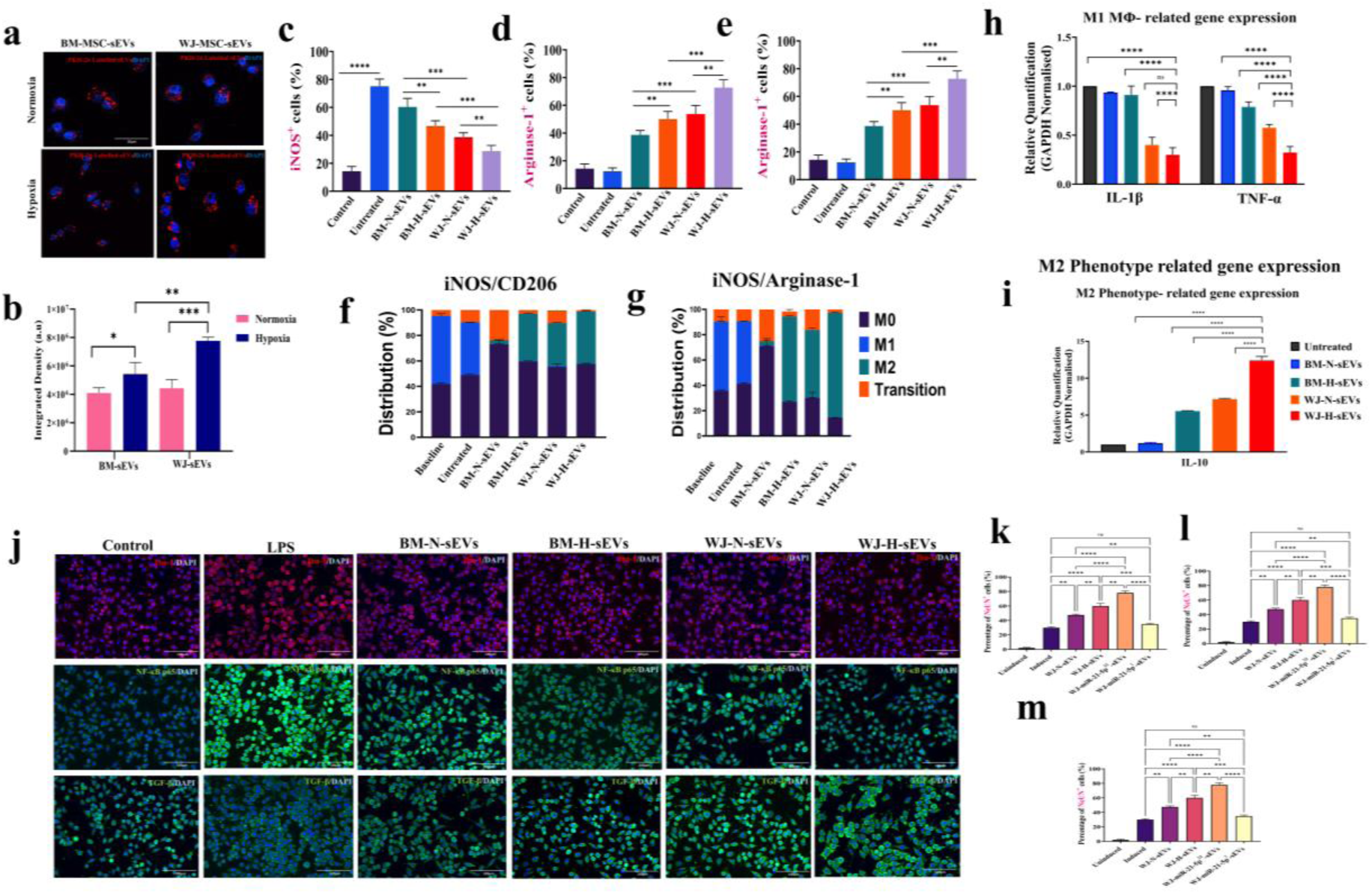
Phenotypic polarization of microglia upon treatment with WJ-H-sEVs. (a) Representative confocal images showing uptake of PKH26-labeled BM- and WJ-MSC-derived sEVs by N9 microglial cells under normoxic and hypoxic conditions. (b) Quantification of sEV internalization by integrated fluorescence density analysis.(c) Percentage of iNOS⁺ microglia following LPS stimulation and treatment with normoxic or hypoxic BM- and WJ-MSC-sEVs. (d,e) Quantification of arginase-1⁺ microglia indicating M2 polarization. (f,g) Distribution of microglial phenotypes (M0, M1, M2, and transitional states) based on iNOS/CD206 and iNOS/Arginase-1 co-expression. (h) Relative mRNA expression of M1-associated cytokines IL-1β and TNF-α measured by qRT-PCR. (i) Relative mRNA expression of the M2-associated cytokine IL-10. (j) Representative immunofluorescence images showing Iba-1, NF-κB p65 nuclear localization, and TGF-β expression in microglia following LPS stimulation and sEV treatment. (k–m) Quantification of NF-κB p65⁺ and TGF-β⁺ cells. Data are presented as mean ± SD (n = 3). Statistical analysis was performed using one-way ANOVA with Bonferroni post hoc test; *p < 0.05, **p < 0.01, ***p < 0.001, ****p < 0.0001. Scale bars: 20 µm. Schematic illustrations were created using BioRender.com.

We next examined microglial polarization by quantifying inducible nitric oxide synthase (iNOS) and arginase-1 expression as markers of M1 and M2 phenotypes, respectively (**Fig. 3c–e**). LPS stimulation robustly increased the proportion of iNOS⁺ microglia, confirming M1 polarization. Treatment with MSC-derived sEVs significantly attenuated this pro-inflammatory response, with WJ-H-sEVs producing the most pronounced reduction in iNOS⁺ cells (p < 0.0001 vs LPS; p < 0.001 vs BM-H-sEVs; p < 0.01 vs WJ-N-sEVs). In parallel, sEV treatment promoted arginase-1⁺ microglia, with WJ-H-sEVs inducing the highest proportion of Arg1⁺ cells among all groups (p < 0.0001 vs BM-N-sEVs and control). These results indicate a robust shift away from classical activation toward a reparative phenotype.

To capture macrophage polarization dynamics more comprehensively, iNOS/CD206 and iNOS/Arginase-1 co-expression analyses were performed (**Fig. 3f,g**). Untreated and LPS-stimulated microglia exhibited a dominant M1 profile, whereas sEV-treated groups showed progressive enrichment of the M2 population. This transition was most prominent in the WJ-H-sEV group, which displayed the lowest M1 fraction and the highest M2 distribution, with minimal intermediate transitional states. These findings suggest that hypoxia-primed WJ-sEVs not only suppress inflammatory activation but actively drive microglia toward a stable, anti-inflammatory phenotype, in line with reports that WJ-MSC-derived EVs exert strong immunosuppressive effects via TLR4/NF-κB interference and induction of alternative activation markers.

Consistent with these phenotypic changes, qRT-PCR analysis revealed that WJ-H-sEVs significantly downregulated M1-associated cytokines IL-1β and TNF-α (p < 0.0001 vs LPS and BM-sEV groups) while markedly upregulating the anti-inflammatory cytokine IL-10 (p < 0.0001 vs all other conditions) (**Fig. 3h,i**). BM-H-sEVs produced moderate suppression of pro-inflammatory genes and induction of IL-10, whereas normoxic sEVs exhibited comparatively weaker effects. These data demonstrate a clear source- and hypoxia-dependent hierarchy in immunomodulatory potency.

To investigate the underlying signaling mechanisms, immunofluorescence analysis of NF-κB p65 nuclear translocation and TGF-β expression was performed following LPS challenge (**Fig. 3j–m**). LPS induced robust nuclear localization of NF-κB p65, indicative of inflammatory pathway activation. Treatment with MSC-derived sEVs significantly attenuated p65 nuclear translocation, with WJ-H-sEVs producing the strongest suppression (p < 0.0001 vs LPS; p < 0.001 vs BM-H-sEVs). These observations align with previous work showing that MSC-EVs downregulate NF-κB activation while boosting IL-10 and TGF-β to promote immune tolerance and neuroprotection.

Collectively, these data demonstrate that hypoxia-primed WJ-MSC-sEVs are preferentially internalized by microglia and exert superior immunomodulatory effects by suppressing NF-κB–driven inflammatory signaling, promoting M2 polarization, and enhancing anti-inflammatory cytokine production. This coordinated regulation of microglial phenotype provides a mechanistic framework linking hypoxia-induced optimization of sEV cargo can lead to restoration of neuroimmune homeostasis in ASD and is consistent with emerging evidence that hypoxia-conditioned MSC exosomes offer enhanced therapeutic benefit in CNS inflammatory disorders.

### Nrf2 Inhibition with ML385 Abolishes WJ-H-sEV-Mediated Antioxidant and Anti-Inflammatory Effects in LPS-Stimulated N9 Microglia

Inhibition or genetic loss of Nrf-2 aggravates oxidative stress, skews microglia towards a pro-inflammatory M1 state, and exacerbates neuroinflammatory injury, whereas MSC-sEVs mediated activation of Nrf-2 has been shown to enhance antioxidant defences and drive anti-inflammatory reprogramming in various CNS disease models. To establish the mechanistic role of WJ-H-sEVs in Nrf2/HO-1 pathway modulation, we employed ML385, a selective Nrf2 inhibitor that disrupts Nrf2-DNA binding at antioxidant response elements (AREs) in LPS-challenged N9 microglia.

Quantitative RT-PCR analysis revealed that LPS stimulation significantly upregulated Nrf2 mRNA expression to 2.2 ± 1.2-fold relative to untreated controls (***p < 0.001 vs. control), reflecting an adaptive transcriptional response to LPS-induced oxidative stress via ROS-mediated Keap1 dissociation. WJ-H-sEV treatment markedly potentiated this to 4.2 ± 0.9-fold (**p < 0.01 vs. LPS alone), consistent with sEV cargo (e.g., miR-21, miR-145) promoting Nrf2 nuclear translocation. However, co-administration of ML385 (LPS + ML385 + WJ-H-sEVs) abrogated this elevation, reducing Nrf2 to 3.5 ± 0.56-fold (**p < 0.01 vs. LPS + WJ-H-sEVs; ns vs. LPS), while ML385 alone yielded 0.75 ± 0.5-fold ((**p < 0.01 vs. control), confirming direct Nrf2 blockade it is abolishing WJ-H-sEVs effect (**Fig. 5a**).

**Figure 5.**
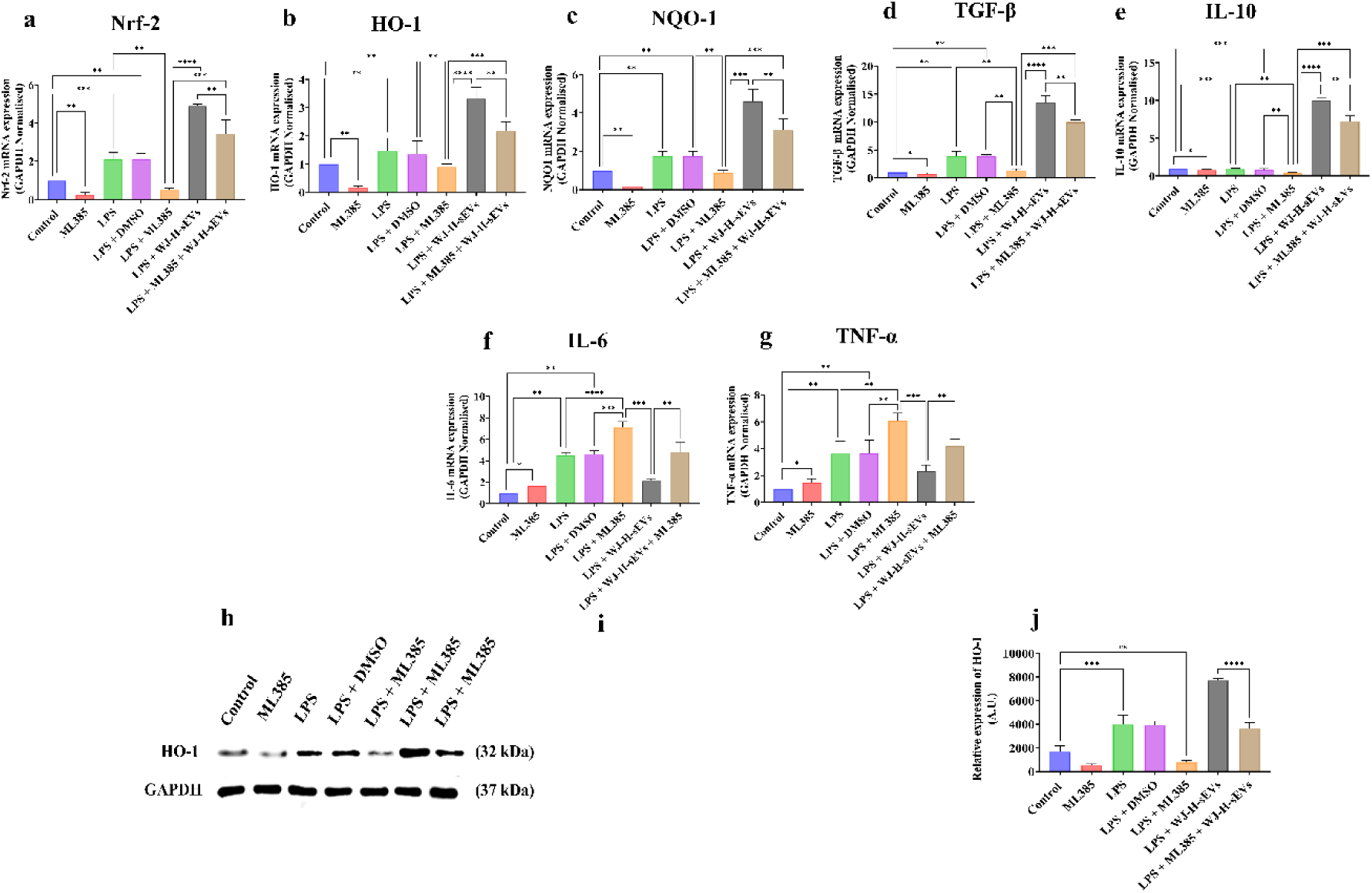
WJ-H-sEVs activate Nrf-2/HO-1 signalling and resolve neuroinflammation in LPS-stimulated N9 microglia in an Nrf2-dependent manner. (a-g) qRT-pCR quantification of relative mRNA expression for Nrf-2 (a), HO-1 (b), NQO1 (c), TGF-β (d), IL-10 (e), IL-6 (f), and TNF-α (g). N9 microglia were pretreated with BAY 11-7082 and then challenged with LPS and followed by WJ-H-sEVs (h) Representative Western Blots showing Nrf2 and HO-1 protein levels in Control, ML385, LPS, LPS + DMSO, LPS+WJ-H-sEVs, and LPS+ML385+WJ-H-sEVs groups, with GAPDH as loading control. (i) Densitometric quantification of Nrf2 and HO-1 bands normalised to GAPDH. Data are presented as mean ± standard deviation (SD, n=3 independent experiments). Statistical analysis was performed using one-way ANOVA followed by Tukey’s post hoc test. ****p<0.0001, ***p<0.001, **p<0.01, *p<0.05, ns = not significant.

This Nrf2-dependence extended robustly to downstream antioxidant proteins. Heme oxygenase-1 (HO-1) increased to 1.56 ± 0.87 fold under LPS (**p < 0.01 vs. control), surged to 3.0 ± 1.04 fold with WJ-H-sEVs (**p < 0.01 vs. LPS) aligning with prior reports of MSC-sEV-induced HO-1 via Nrf2-ARE signaling in neuroinflammatory models and reverted to 1.9 ± 0.2-fold upon ML385 addition (**p < 0.01 vs. LPS + WJ-H-sEVs; ns vs. LPS), demonstrating essential Nrf2 mediation by WJ-H-sEVs (**Fig. 5b**). NAD(P)H quinone dehydrogenase 1 (NQO1) mirrored this profile, LPS induced 2.0 ± 0.2-fold (**p < 0.01 vs. control), WJ-H-sEVs elevated to 4.2 ± 1.2-fold (**p < 0.01 vs. LPS), and ML385 suppressed to 3.1 ± 0.82-fold (**p < 0.01 vs. LPS + WJ-H-sEVs; ns vs. LPS), underscoring coordinated Nrf2 orchestration of phase II detoxification enzymes (**Fig. 5c**).

WJ-H-sEV-driven anti-inflammatory reprogramming was similarly Nrf2-dependent. Transforming growth factor-β (TGF-β), minimally altered by LPS (1.1 ± 0.1-fold, ns vs. control), exhibited dramatic induction to 13.2 ± 1.2-fold by WJ-H-sEVs (***p < 0.001 vs. LPS) a response recapitulating sEV-mediated M2 polarization in microglia and collapsed to 1.3 ± 0.1-fold with ML385 (**p < 0.001 vs. LPS + WJ-H-sEVs; ns vs. LPS), linking Nrf2 to immunoregulatory cytokine homeostasis (**Fig. 5d**).

IL-10 exhibited similar dynamics, the expression remains unchanged by LPS (1.0 ± 0.1-fold, ns), elevated to 9.5 ± 0.8-fold (WJ-H-sEVs, ***p < 0.001 vs. LPS), and reversed to 1.2 ± 0.1-fold (ML385, **p < 0.01 vs. LPS + WJ-H-sEVs), affirming Nrf2 as a requisite hub for sEV anti-inflammatory efficacy (**Fig. 5e**).

Concomitantly, WJ-H-sEV suppression of pro-inflammatory cytokines required Nrf2 signaling as evident by the results. LPS drove interleukin-6 (IL-6) to 4.5 ± 0.4-fold (**p < 0.01 vs. control); WJ-H-sEVs attenuated to 2.45 ± 0.3-fold (*p < 0.05 vs. LPS), but ML385 restored levels to 4.2 ± 0.4-fold (***p < 0.001 vs. LPS + WJ-H-sEVs; ns vs. LPS), revealing Nrf2-mediated repression of NF-κB-driven IL-6 (**Fig. 5f)**. Tumor necrosis factor-α (TNF-α) peaked at 4.5 ± 0.4-fold (LPS, ****p < 0.0001 vs. control), declined to 2.0 ± 0.2-fold (WJ-H-sEVs, ***p < 0.001 vs. LPS), and rebounded to 4.1 ± 0.3-fold (ML385, ***p < 0.001 vs. LPS + WJ-H-sEVs; ns vs. LPS), confirming Nrf2 as the upstream gatekeeper of sEV immunomodulation (**Fig. 5g**).

Western blot/Protein analysis via WB post 48 h all post-treatment LPS (**Fig. 5h**) corroborated mRNA trends, with densitometric quantification showing Nrf2 (**Fig.5i**), and HO-1 (**Fig.5j**) protein elevations in LPS + WJ-H-sEVs versus LPS, abolished in ML385-co-treated samples thus affirming Nrf2 reliance of WJ-H-sEVs.

Collectively, these data unequivocally demonstrate that WJ-H-sEV neuroprotective effects are Nrf2 dependent as evidenced by comprehensive ML385 ab. This establishes Nrf2 as the primary effector of WJ-H-sEV action which has the potential to restore neuronal homeostasis and mitigate ASD core symptoms as documented by previous studies where inflammatory and oxidative markers correlated with the severity of ASD and also the results establish the mechanism of action of WJ-H-sEVs.

### NF-κB Inhibition Potentiates WJ-H-sEV–Mediated Antioxidant and Anti-Inflammatory Effects in LPS-Stimulated N9 Microglia

After establishing the dependence of WJ-H-sEV-mediated effects on Nrf2 activation using ML385, we next evaluated whether pharmacological inhibition of NF-κB with BAY 11-7082 restores or enhances WJ-H-sEV-driven antioxidant function under LPS-induced inflammatory stress in N9 microglia.

Quantitative RT-PCR analysis revealed that LPS stimulation significantly upregulated Nrf2 expression to 2.2-fold relative to untreated control cells (***p < 0.001 vs. control), consistent with compensatory Nrf2 activation in response to oxidative stress in microglia. Treatment with WJ-H-sEVs in LPS-stimulated N9 microglia further elevated Nrf2 levels to 3.2-fold over control (**p < 0.01 vs. LPS alone), reflecting the inherent antioxidant potential WJ-H-sEVs. Co-treatment with BAY 11-7082 (LPS + BAY + WJ-H-sEVs) increased Nrf2 to 3.8-fold (***p < 0.001 vs. LPS), supporting established negative cross-regulation between NF-κB and Nrf2, where NF-κB inhibition prevents Nrf2 nuclear exclusion or competitive DNA binding (**Fig. 6a**).

**Figure 6.**
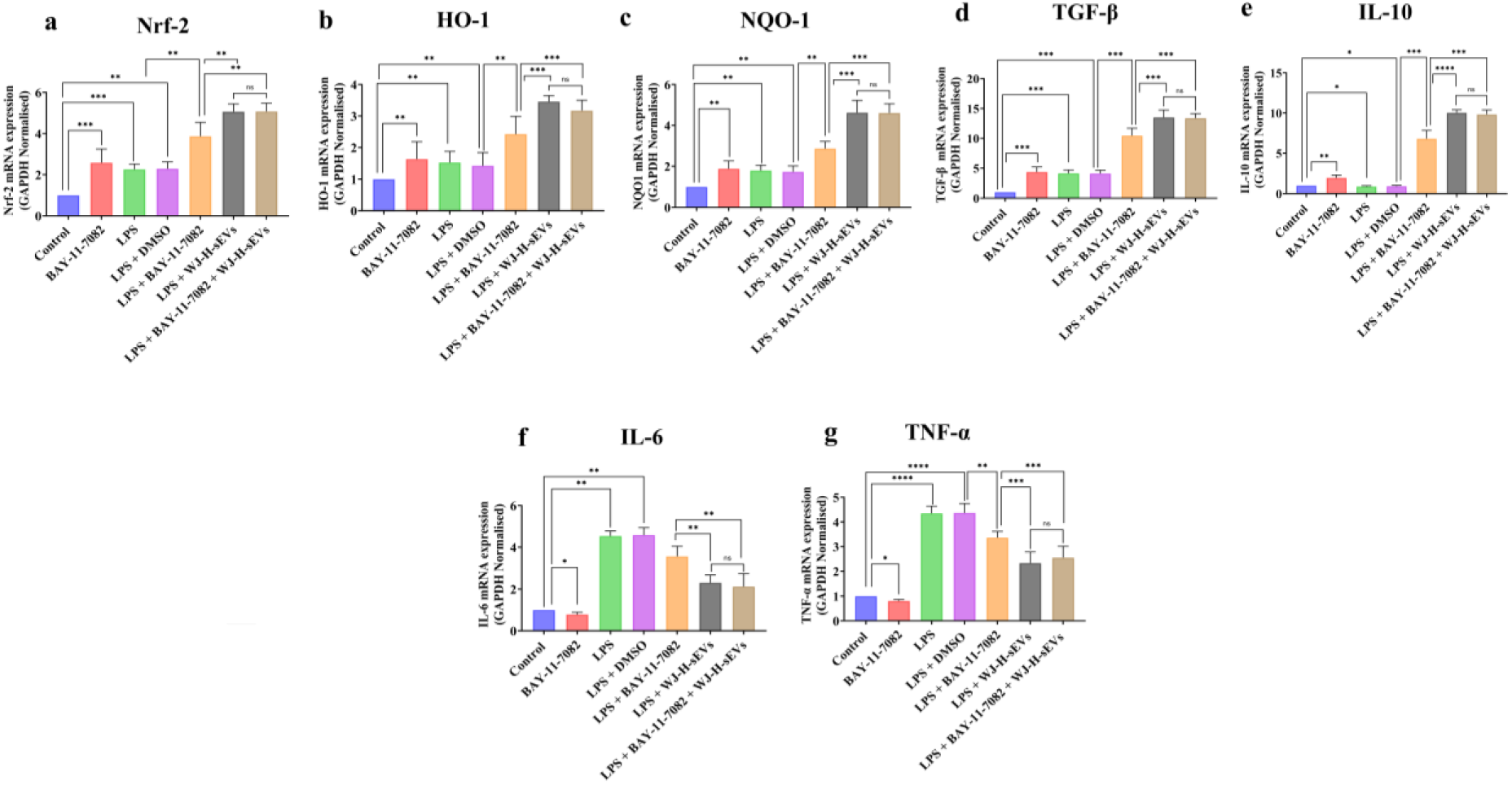
NF-κB inhibition potentiates WJ-H-sEV-mediated antioxidant gene upregulation and cytokine modulation in LPS-stimulated N9 microglia. Quantitative real-time PCR analysis of relative mRNA expression (fold change normalized to control) for (a) Nrf2, (b) HO-1, (c) NQO1, (d) TGF-β, (e) IL-10, (f) IL-6, and (g) TNF-α. N9 microglia were pretreated with BAY 11-7082 and then challenged with LPS and followed by WJ-H-sEVs. Data are presented as mean ± standard deviation (SD, n=3 independent experiments). Statistical analysis was performed using one-way ANOVA followed by Tukey’s post hoc test. ****p<0.0001, ***p<0.001, **p<0.01, *p<0.05, ns = not significant

HO-1 expression rose to 2.0-fold with LPS (**p < 0.01 vs. control), reached 3.0-fold upon WJ-H-sEV addition (**p < 0.01 vs. LPS), and trended higher to ∼3.4-fold with BAY co-treatment (***p < 0.001 vs. LPS; ns vs. LPS + WJ-H-sEVs), indicating that WJ-H-sEVs achieve near-maximal HO-1 induction independently of NF-κB blockade (**Fig. 6b**). NQO1 followed suit, with LPS inducing ∼2.0-fold (**p < 0.01 vs. control), WJ-H-sEVs elevating to ∼3.0-fold (**p < 0.01 vs. LPS), and BAY combination yielding ∼3.2-fold (***p < 0.001 vs. LPS; ns vs. LPS + WJ-H-sEVs) (**Fig. 6c**).

In anti-inflammatory cytokines TGF-beta and IL-10, and we found that LPS minimally affected TGF-β (1.1-fold, ns vs. control), while WJ-H-sEVs robustly induced 13.2-fold (**antinflammatory action of WJ-H-sEVs expression**) (***p < 0.001 vs. LPS); BAY co-treatment also showed increased to ∼12.8-fold (***p < 0.001 vs. LPS; ns vs. LPS + WJ-H-sEVs), highlighting sustained but non-significantly enhanced resolution signaling (**Fig. 6d**). IL-10 WJ-H-sEV stimulation to 9.5-fold (***p < 0.001 vs. LPS), and BAY addition to 9.34-fold (***p < 0.001 vs. LPS; ns vs. LPS + WJ-H-sEVs), supporting cooperative yet plateaued anti-inflammatory amplification (**Fig. 6e**).

Pro-inflammatory markers were effectively curtailed. LPS induced IL-6 to ∼4.5-fold (**p < 0.01 vs. control), reduced by WJ-H-sEVs to 2.45-fold (*p < 0.05 vs. LPS), with BAY co-treatment lowering to 2.31-fold (**p < 0.01 vs. LPS; *p < 0.05 vs. LPS + WJ-H-sEVs), demonstrating additive suppression here (**Fig. 6f**). TNF-α spiked to ∼4.5-fold under LPS (****p < 0.0001 vs. control), fell to 2.0-fold with WJ-H-sEVs (***p < 0.001 vs. LPS), and remained at 2.3-fold with BAY (***p < 0.001 vs. LPS; ns vs. LPS + WJ-H-sEVs), evidencing maximal sEV-mediated restraint unaffected by further NF-κB inhibition (**Fig. 6g**).

These findings align with prior reports of Nrf2-NF-κB antagonism in microglia, where NF-κB inhibitors like BAY 11-7082 alleviate suppression of Nrf2-driven antioxidative genes (e.g., HO-1, NQO1) and cytokines, thereby potentiating MSC-sEV therapeutics for neuroinflammatory disorders such as autism spectrum disorder as the underlying pathology is oxinflammation and MSC-sEVs as documented in opther indications as well can effectively control both oxidative stress and inflammation and the results of this study also demonstrate the same.

These observations substantiate the antagonistic interplay between the Nrf2 and NF-κB pathways in microglia, whereby NF-κB inhibitors such as BAY 11-7082 mitigate transcriptional repression of Nrf2 target genes, including HO-1 and NQO1, thereby augmenting antioxidative gene expression and restoring cytokine homeostasis. By re-establishing redox–immune balance through coordinated modulation of these antagonistic pathways, WJ-H-sEVs represent a mechanistically rational biological strategy to interrupt the self-sustaining oxinflammatory cascades implicated in ASD and address the existing therapeutic gap.

### Intranasal route facilitates brain-specific localization of WJ-H-sEVs compared to intravenous injection

To evaluate the spatiotemporal biodistribution of WJ-H-sEVs, as outlined in the schematic (**Fig. 5a**), PKH26-labeled WJ-H-sEVs (100 μg) were delivered via intranasal and intravenous routes to normal and VPA-exposed pups at PND-42, followed by *ex-vivo* organ biodistribution analysis at 3 h, 6 h, and 12 h post-administration.

*Ex vivo* fluorescence imaging of harvested organs demonstrated prominent accumulation of PKH26-labeled WJ-H-sEVs in the brain following intranasal delivery, with comparatively lower signal intensities detected in peripheral organs including liver, spleen, lung, heart, and kidney. Robust fluorescent signals of PKH-26 labelled sEVs were detected in the brain as early as 3 h post-delivery, with signal intensity progressively increasing at 6 h and persisting up to 12 h (**Fig. 5b**). However, PBS-treated animals did not show detectable fluorescence, confirming signal specificity and the absence of autofluorescence.

Quantitative analysis of radiant efficiency 3 h post-administration demonstrated (**Fig. 5c**) mean brain radiant efficiency in PBS-treated animals remained low (1.2 × 10⁷ ± 0.3 × 10⁷ p/s/cm²/sr). In contrast, intranasal WJ-H-sEV treatment resulted in significantly higher brain signals in control mice (3.4 × 10⁷ ± 0.6 × 10⁷ p/s/cm²/sr) and VPA mice (3.8 × 10⁷ ± 0.7 × 10⁷ p/s/cm²/sr) (p < 0.01 vs PBS). No significant increase was observed in liver, spleen, lung, heart, or kidney (ns). By 6h, (**Fig. 5d**) brain fluorescence further increased, reaching 5.2 × 10⁷ ± 0.8 × 10⁷ p/s/cm²/sr in control mice and 5.9 × 10⁷ ± 0.9 × 10⁷ p/s/cm²/sr in VPA mice, which was significantly higher than all peripheral organs (p < 0.001). Peripheral organ signals remained low and statistically indistinguishable from PBS control. At 12h post administration (**Fig. 5e**), brain-associated radiant efficiency was the highest (7.1 × 10⁷ ± 1.0 × 10⁷ p/s/cm²/sr) in control mice and (8.3 × 10⁷ ± 1.2 × 10⁷ p/s/cm²/sr) in VPA mice. These values were significantly greater than fluorescence measured in liver, spleen, lung, kidney, and heart (p < 0.0001), confirming sustained and preferential brain retention of intranasally delivered WJ-H-sEVs.

Time-course analysis of brain fluorescence (**Fig. 5d**) corroborated these findings, showing a progressive increase in radiant efficiency from 3 h to 12 h in both groups. In control mice, brain signal increased from (3.4 × 10⁷ ± 0.6 × 10⁷ p/s/cm²/sr) at 3 h to (7.1 × 10⁷ ± 1.0 × 10⁷ p/s/cm²/sr) at 12 h, while VPA mice exhibited a rise from (3.8 × 10⁷ ± 0.7 × 10⁷ p/s/cm²/sr) to (8.3 × 10⁷ ± 1.2 × 10⁷ p/s/cm²/sr) over the same period (p < 0.01 at 3 h; p < 0.001 at 6 h; p < 0.0001 at 12 h vs PBS). Together, these data align with previously published reports^63,64^ and demonstrate that intranasal administration enables efficient, selective, and sustained delivery of WJ-H-sEVs to the brain, with minimal exposure to peripheral organs, supporting this non-invasive route as a clinically translatable strategy for sEV-based neurotherapeutic interventions in ASD.

### WJ-H-sEVs delivered through the intranasal route enhance recognition memory while alleviating locomotor and exploration deficits in VPA model mice

Prenatal VPA exposure is a well-validated model of ASD, recapitulating early neurodevelopmental delays, increased anxiety, reduced exploratory drive, and cognitive impairments that parallel clinical observations in autistic children. Consistent with previous reports, VPA-exposed offspring in this study was screened to from PND-9-21 as schematized in Fig.1 to determine early behavioural features. VPA offsprings showed significantly lower body weight on PND-21 (p < 0.05). A 2-day delay in eye opening was found in the VPA group (Fig.S). While control pups had both eyes opened by PND-14 (p < 0.01), it was not until PND16 that VPA offsprings did so. These results indicate delayed maturation, growth and behavioural development in VPA offsprings.

To determine whether WJ-H-sEVs can ameliorate ASD-like behavioral abnormalities, 100 µg of WJ-H-sEVs were administered intranasally once every 4 weeks starting at PND-42, followed by open field test (OFT) and novel object recognition testing (NORT) as outlines in the schematic outline (**Fig.6a**). In the OFT (**Fig.6b**), similar to previous reports of other ASD model, VPA offspring showed marked anxiety-like behaviour and reduced exploratory drive compared with controls, as reflected by a significantly decreased percentage of time spent in the central zone (Control: 42.31 ± 3.87%, VPA: 19.62 ± 3.11%, p < 0.01) and a corresponding increase in time spent in the peripheral zone (Control: 57.69 ± 3.87%, VPA: 80.38 ± 3.11%, p < 0.01) (**Fig. 6c–e**). The open-field exploration index (OFEI) was likewise reduced in VPA mice (Control: 0.71 ± 0.05, VPA: 0.32 ± 0.04, p < 0.001), confirming exploration deficits (**Fig.6f**). Intranasal WJ-H-sEV treatment significantly increased centre time in VPA mice (VPA + WJ-H-sEVs: 36.84 ± 4.02%, p < 0.05 vs VPA, ns vs Control) and reduced peripheral time (VPA + WJ-H-sEVs: 63.16 ± 4.02%, p < 0.05 vs VPA), resulting in partial normalisation of OFEI (VPA + WJ-H-sEVs: 0.63 ± 0.06, p < 0.05 vs VPA, ns vs Control (**Fig. 6c–f**). Notably, Control + WJ-H-sEVs mice did not differ from untreated controls in centre time, peripheral time, or OFEI (all ns), indicating that WJ-H-sEVs did not adversely affect exploration or anxiety-related behaviour in healthy animals and supporting their behavioural safety profile, however the results suggest an improvement in exploration deficits and anxiety-like behavior within the WJ-H-sEVs treated group.

In the NORT (**Fig.6g**), VPA offspring exhibited a clear deficit in recognition memory compared with controls, spending significantly less time exploring the novel object (Control: 70 ± 4.45%, VPA: 35 ± 1.25%, p < 0.001) and more time with the familiar object (Control: 30 ± 2.45%, VPA: 65 ± 4%, p < 0.01; (**Fig. 6h–j**). This shift in exploration pattern was reflected in a reduced discrimination index in VPA mice (Control: ∼0.85 ± 0.05, VPA: ∼0.40 ± 0.05, p < 0.001), confirming impaired recognition memory (**Fig. 6k**). Intranasal WJ-H-sEV treatment robustly rescued these deficits: VPA + WJ-H-sEVs mice spent significantly more time exploring the novel object (∼60 ± 4%, p < 0.01 vs VPA, ns vs Control) and less time with the familiar object (∼40 ± 4%, p < 0.05 vs VPA), with discrimination index values restored to near-control levels (∼0.75 ± 0.04, p < 0.01 vs VPA, ns vs Control (**Fig. 6h–k**). Importantly, Control + WJ-H-sEVs animals did not differ from untreated controls in time spent with the novel or familiar object or in discrimination index (all ns), indicating that WJ-H-sEVs do not impair cognitive performance in control mice.

Collectively, these data show that intranasal WJ-H-sEVs reverse VPA-induced anxiety-like behaviour, exploration deficits, and recognition-memory impairments, with no change in behaviour of control mice thereby supporting a favourable safety profile for this intervention. This behavioural rescue is consistent with prior work in ASD^63,65,66^ and related neurodevelopmental^67–71^/neurodegenerative models^72–78^. MSCs and their derived sEVs can restore excitatory–inhibitory balance^79–81^, enhance dendritic spine density^82,83^ and neurite growth^84,85^, promote neural progenitor proliferation^86,87^, and normalise long-term potentiation^78,86^ in hippocampal and cortical circuits by delivering neuroprotective miRNAs^43,52,77,88–93^ that modulate PTEN/PI3K–AKT^43^, NF-κB^49,93^, and BDNF–ERK–CREB^76^ signalling and regulate gene sets involved in synaptic plasticity and neuronal function^72–74,78,88^. Thus, hypoxia-primed WJ-sEVs are likely to act by reprogramming dysregulated signalling networks and synaptic plasticity within hippocampal–prefrontal circuits, thereby ameliorating ASD-like phenotypes in the VPA model.

### Amelioration of spatial and working memory impairments by WJ-H-sEVs administration in VPA model mice

Next, we evaluated the spatial learning and long term working memory using the Morris Water Maze test (**Fig. 7a**). A six-day MWM experiment was conducted for each group of mice. On day 6, VPA offspring required 38.67 ± 6.8 s to reach the platform versus 20.2 ± 4.8 s in controls (p < 0.01), consistent with previous studies. In contrast, WJ-H-sEV-treated VPA mice showed substantially shorter escape latencies of 23.0 ± 6.7 s on day 6, significantly improved compared with untreated VPA animals (p < 0.01) and not different from control (21.9 ± 5.2 s, ns) or WJ-H-sEV-treated control groups (22.3 ± 7.4 s, ns), indicating complete normalisation of spatial acquisition. Control + WJ-H-sEVs mice performed similarly to untreated controls at all time points (ns), supporting the cognitive safety of WJ-H-sEVs in healthy animals (**Fig. 7b–d**).

**Figure 7.**
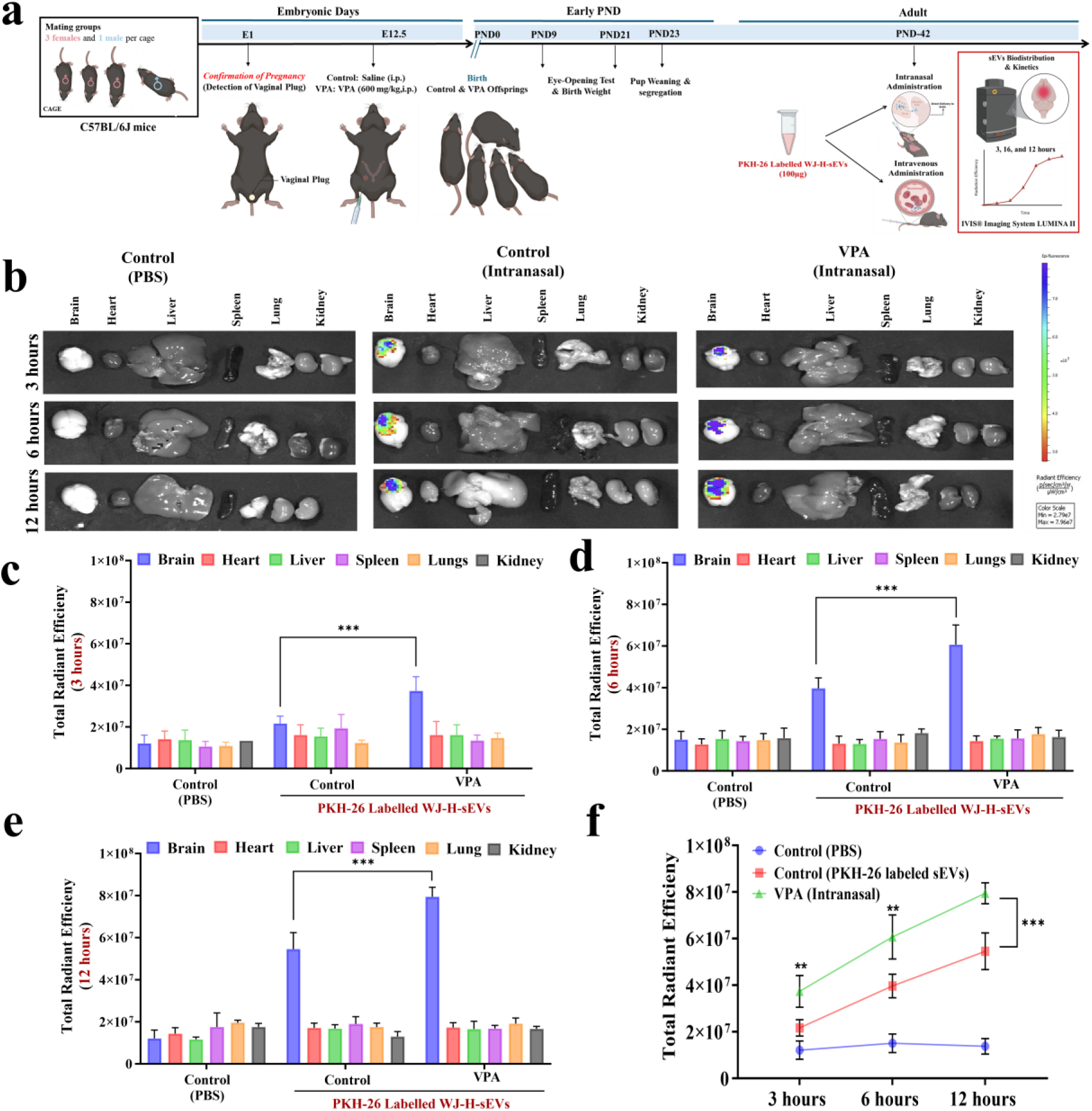
Intranasal administration reveals preferential brain targeting of WJ-H-sEVs in *ex-vivo* biodistribution analysis. (a) Schematic illustration of the experimental design depicting intranasal and intravenous administration of PKH26-labeled WJ-H-sEVs (100 µg) to control and VPA-exposed offspring at PND-42, followed by ex vivo biodistribution analysis at 3 hours, 6 hours, and 12 hours post-administration. (b) Representative ex vivo IVIS® fluorescence images of harvested organs (brain, liver, spleen, lung, heart, and kidney) from control and VPA animals following intranasal administration of PKH26-labeled WJ-H-sEVs. (c–e) Quantitative ex vivo analysis of PKH26 fluorescence expressed as radiant efficiency (p/s/cm²/sr) in the brain and peripheral organs at 3 hours (c), 6 hours (d), and 12 hours (e) post-administration following intranasal delivery of WJ-H-sEVs. (f) Time-course analysis of brain-associated fluorescence showing changes in radiant efficiency across the time points (3h, 6h and 12h) following intranasal administration in control and VPA-exposed mice. Data are presented as mean ± standard deviation (SD). Statistical analysis was performed using one-way ANOVA ollowed by Tukey’s post hoc test. ****p<0.0001,***p<0.001,,**p<0.01, *p < 0.05, ns = not significant. (Schematic created with BioRender.com)

Probe-trial measures confirmed a pronounced deficit in long-term working memory in VPA offspring and its rescue by WJ-H-sEVs (**Fig. 7e-f**). VPA mice exhibited a severely reduced number of platform-site crossings (2.67 ± 5.90 vs. 10.89 ± 7.58 in controls, p < 0.01) and a prolonged latency to the first target-site cross-over (40.67 ± 6.98 s vs. 18.45 ± 4.67 s in controls, p < 0.01). WJ-H-sEV treatment increased platform crossings in VPA mice to 11.45 ± 4.8 s and reduced probe time to 20.09 ± 5.8 s, both significantly improved relative to untreated VPA animals (p < 0.01) and indistinguishable from control groups (ns), indicating restored memory for the platform location.

Consistent with these indices, VPA mice spent only ∼12 ± 3% of total time and a similar fraction of swim distance in the target quadrant, significantly less than controls (35.56 ± 5.87%, p < 0.01) (**Fig. 7g-h**). WJ-H-sEV-treated VPA mice showed a marked increase in target-quadrant occupancy and distance to 32.78 ± 5.03%, statistically indistinguishable from controls (ns) and significantly higher than untreated VPA animals (p < 0.01).

Together with the OFT and NORT findings, these data demonstrate that intranasally delivered WJ-H-sEVs not only alleviate anxiety-like behaviour and recognition-memory deficits but also effectively rescue VPA-induced impairments in spatial learning and long-term working memory. This benefit is likely mediated by neuroprotective miRNAs^44,48,77,91,92^ with their multimodal actions^94^ on hippocampal–prefrontal cortex circuit, particularly CA1/CA3^86^ and medial-prefrontal pyramidal neurons^81^ and their local interneuron networks^95^. MSC-derived sEVs have been shown to protect these neurons from apoptosis^85^ and to stimulate neurogenesis^86,87^ in the dentate gyrus and other neurogenic niches^67^, leading to improved cognitive performance in diverse neurological models. Overall, such multidomain cognitive benefits of WJ-H-sEVs are highly clinically desirable in the context of ASD.

### WJ-H-sEV treatment attenuates neuroinflammation and restores synaptic integrity in the brain of VPA-induced ASD mice

OxInflammation in ASD is closely associated with synaptic dysfunction and behavioral impairments in both patients and experimental models. To determine whether WJ-H-sEVs modulate neuroinflammation in ASD, we evaluated region-specific (hippocampus and prefrontal cortex) glial activation, systemic cytokine profiles, and synaptic protein expression following intranasal administration of WJ-H-sEVs in the VPA mouse model (**Fig.8**).

**Fig. 8.**
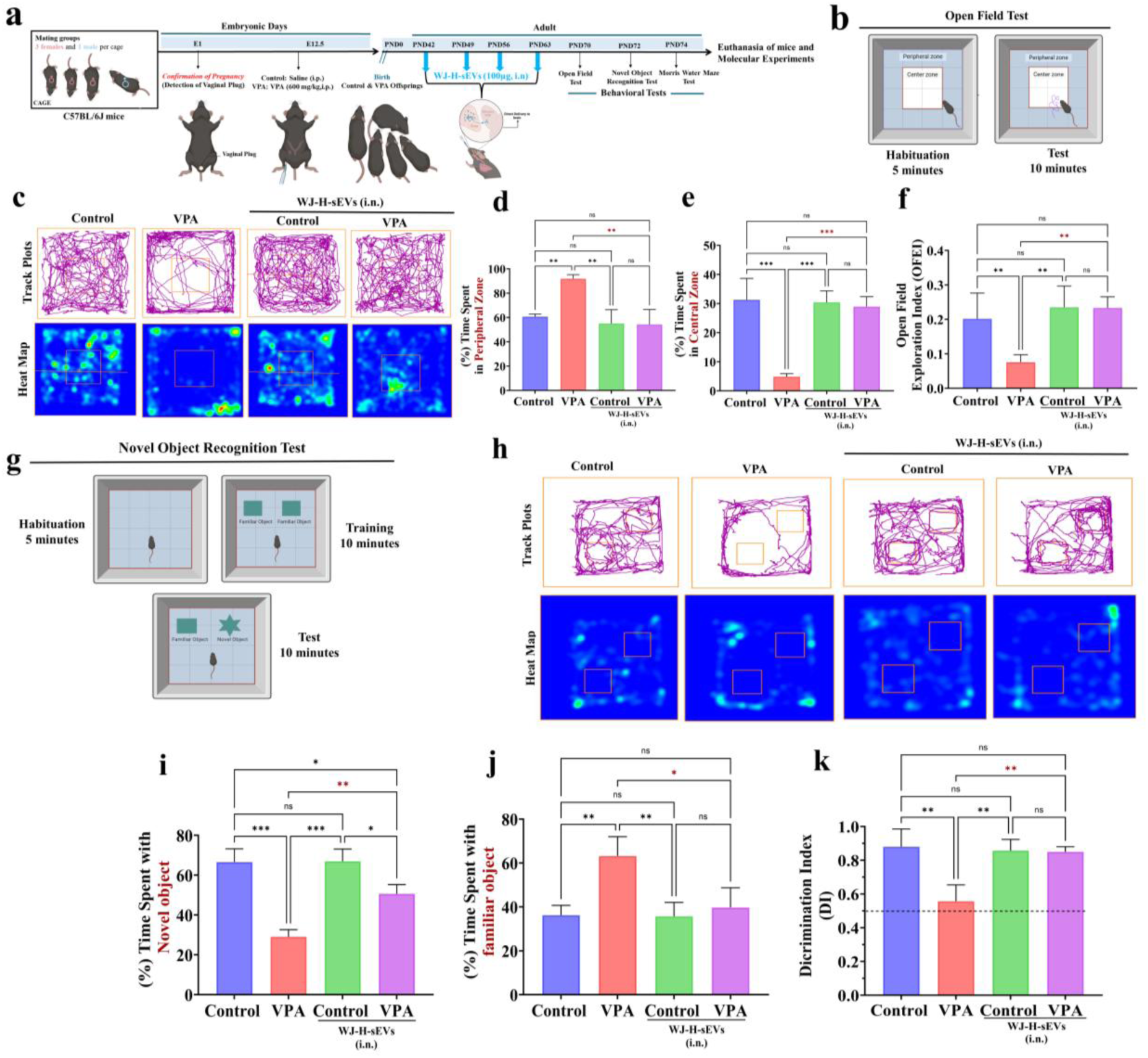
Therapeutic Effects of WJ-H-sEVs on Memory, Locomotor, and Exploration Deficits in the VPA offspring. (A) Schematic diagram illustrating the experimental procedure of sEVs administration and timeline and subsequent behavioural studies conducted. (B) Schematic illustration of the open field area. (C) Representative track and heatmap images from the tested mice (C) Quantification of the time spent in the central zone. (D) Quantification of the time spent in the peripheral zone. (E) OFEI of mice in the open field tasks. (F) Experimental design of novel object recognition test (G) Representative Track plots and Heat Maps from the tested mice in the novel object recognition test (H) Time spent with the with the familiar object (I) Time spent with the novel object percentage (J) DI of each group of mice. All data are presented as the mean ± SD,n=3. Two-way ANOVA followed by Bonferroni post hoc test. ****p<0.0001,***p<0.001,,**p<0.01, *p < 0.05, ns p ≥0.05. Abbreviations: DI, Discrimination index; VPA, valproic acid. (Schematic created with BioRender.com)

PFC function VPA-exposed mice exhibited a pronounced neuroinflammatory phenotype characterized by robust astrocytic and microglial activation in the prefrontal cortex and hippocampus in line with prior reports. Quantitative immunofluorescence analysis revealed pronounced reactive gliosis in the prefrontal cortex (PFC) of VPA-exposed mice. VPA mice exhibited significantly elevated GFAP-positive astrocytes and Iba-1-positive microglia compared with controls (*p* < 0.01 for both markers; **Fig. 8a–e**), indicating sustained astrocytic hypertrophy and microglial activation. These findings are consistent with post-mortem ASD studies and VPA models linking cortical gliosis to disrupted excitatory–inhibitory balance and impaired synaptic signaling. Intranasal WJ-H-sEV administration significantly reduced GFAP immunoreactivity (*p* < 0.05 vs. VPA) and Iba-1 immunoreactivity (*p* < 0.01 vs. VPA), restoring glial marker levels to those comparable with control animals receiving WJ-H-sEVs (ns), suggesting effective suppression of pathological gliosis rather than nonspecific glial loss.

A similar neuroinflammatory profile was observed in the hippocampus, a region critical for learning, memory, and emotional regulation. VPA exposure induced robust astrocytic and microglial activation, reflected by significantly increased GFAP and Iba-1 staining relative to controls (*p* < 0.01 for both markers; **Fig. 8i–j**). Hippocampal glial activation has been widely implicated in ASD-associated synaptic plasticity deficits and cognitive dysfunction. WJ-H-sEV treatment significantly attenuated both astrocytic and microglial activation in the hippocampus (*p* < 0.01 vs. VPA for GFAP and Iba-1), normalizing glial reactivity toward control levels (ns), indicating a consistent anti-inflammatory effect across cortico-limbic regions.

Western blot analysis of whole-brain lysates corroborated the immunofluorescence findings (**Fig. 8k–q**). VPA mice exhibited significant upregulation of GFAP and Iba-1 protein expression compared with controls (*p* < 0.01 for both), confirming neuroinflammatory activation. WJ-H-sEV treatment significantly reduced GFAP (*p* < 0.05 vs. VPA) and Iba-1 (*p* < 0.01 vs. VPA) protein levels, with no significant difference compared with control mice receiving WJ-H-sEVs, reinforcing the concordance between regional histological and biochemical outcomes.

Consistent with central neuroinflammatory alterations, VPA exposure also induced a systemic pro-inflammatory cytokine shift, characterized by elevated serum IL-6 and reduced IL-10 levels compared with controls (*p* < 0.01 for both; **Fig. 8l–m**). Elevated IL-6 has been implicated in ASD severity and synaptic dysfunction via sustained STAT3 and NF-κB signaling, whereas reduced IL-10 reflects impaired anti-inflammatory resolution. WJ-H-sEV treatment significantly reduced the elevated serum IL-6 levels observed in VPA mice (*p* < 0.05 vs. VPA) while concomitantly restoring anti-inflammatory IL-10 levels (*p* < 0.01 vs. VPA; **Fig. 8l–m**), consistent with prior reports demonstrating that MSC-derived sEVs suppress pro-inflammatory cytokine release while enhancing anti-inflammatory signaling through modulation of macrophage/microglial polarization and inhibition of NF-κB dependent transcription.

Inflammation-induced activation of NF-κB and JAK-STAT signaling, together with elevated reactive oxygen species (ROS), exacerbates synaptic loss by suppressing PSD-95 and synaptophysin expression, impairing NMDA receptor function, and disrupting dendritic spine morphogenesis. Given the close coupling between neuroinflammation and synaptic stability, synaptic protein expression was therefore examined (**Fig. 8j)**. VPA exposure resulted in a significant reduction in the postsynaptic scaffolding protein PSD-95 and the presynaptic marker synaptophysin (*p* < 0.01 vs. control). Notably, WJ-H-sEV treatment significantly restored the expression of both synaptic markers (*p* < 0.01 vs. VPA), indicating recovery of synaptic integrity in affected brain regions (Hippocampus and Prefrontal Cortex) (**Fig. 8m-n)**.

Collectively, these findings demonstrate that intranasal WJ-H-sEVs mitigate VPA-induced glial activation, normalize systemic inflammatory profiles, and rescue synaptic protein expression in the prefrontal cortex and hippocampus of the ASD mice model. By targeting inflammation-driven synaptic pathology, WJ-H-sEVs address core neuropathological mechanisms underlying ASD and highlight their therapeutic potential for ameliorating OxInflammation-associated synaptic dysfunction.

### WJ-H-sEVs activated the Nrf2 antioxidant pathway and concomitantly suppressed the JAK2/STAT3 and NF-κB inflammatory axes

Further, we investigated the molecular basis of the neuroprotective effects mediated by WJ-H-sEVs in the VPA brain to determine whether the observed behavioral and neuroinflammation and oxidtiave tress improvements reflect correction of core pathogenic signaling abnormalities.

Immunoblot and quantitative analysis (**Fig.9**) reveal that VPA exposure leads to significant upregulation of RelA, NFkB-1, Iba-1, and GFAP, indicating heightened neuroinflammation and glial activation, while Nrf-2 levels are relatively suppressed (**Fig.9a-c**). Upon intranasal administration of WJ-H-sEVs, there is a pronounced reversal of these inflammatory markers, with NF-kB pathway components significantly reduced, and Nrf-2 expression restored, underscoring the anti-inflammatory and neuroprotective properties of the sEVs *in vivo* (**Fig.9a-c**).

**Fig. 9.**
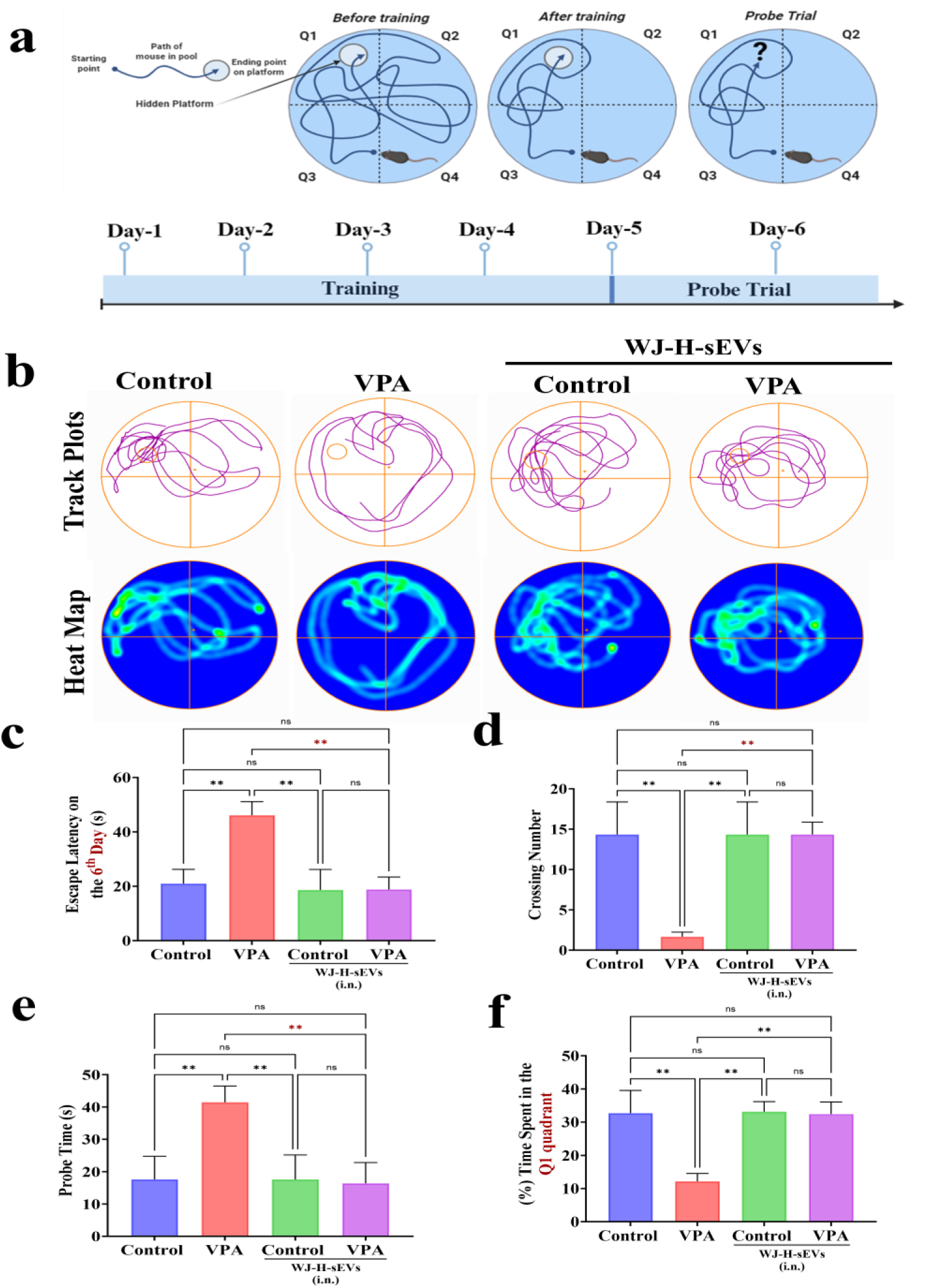
Restoration of spatial memory and long-term working memory by WJ-H-sEVs in the Morris Water Maze Test. (a) Illustration of the morris water maze and the timeline of spatial acquisition and probe trial sessions (b) Representative swimming paths of the mice, receiving sham, treatment and control during the probe trial (c) The escape latencies on the 7^th^ day of the spatial acquisition session. (e-f) The average crossing number over the platform-site and the latency of the first target-site cross-over (probe time) during the probe trial. (g-h) The percentage of time spent and distance travelled in the target quadrant during the probe trial. n=4 for each group. Data were presented as mean ± SD. ****p<0.0001,***p<0.001,,**p<0.01, *p < 0.05, ns = not significant. Schematic created with BioRender.com)

**Fig. 10.**
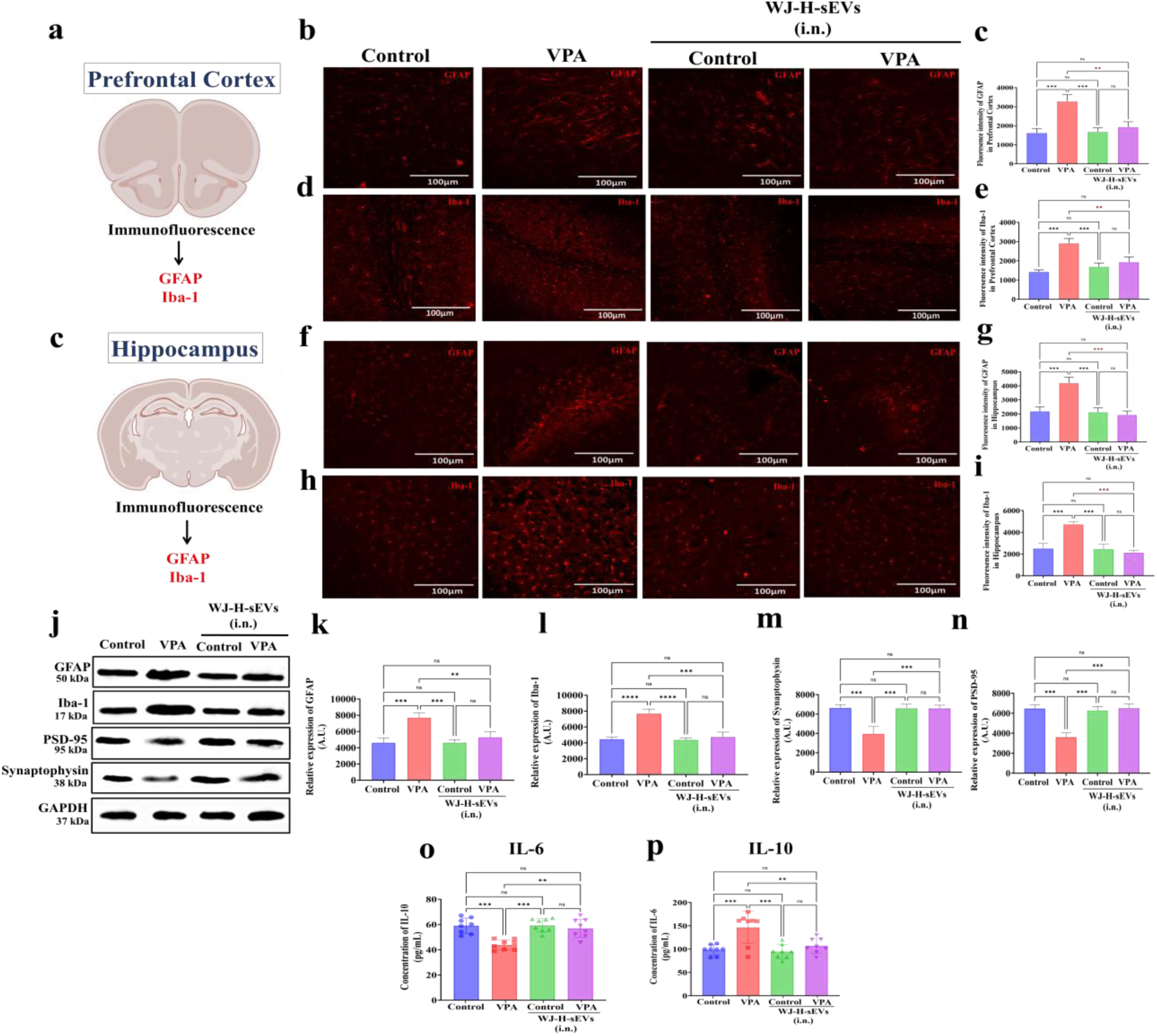
Restoration of neuroinflammation by WJ-H-sEVs in prefrontal cortex and hippocampus of VPA ASD model mice. (a) Schematic of prefrontal cortex region analyzed. (b) Representative immunofluorescence images of GFAP (red fluorescence) and (c) quantitative analyses of GFAP mean fluorescence intensity in prefrontal cortex. (d) Representative immunofluorescence images of Iba-1 (red fluorescence) and (e) quantitative analyses of Iba-1 mean fluorescence intensity in prefrontal cortex. (f) Schematic of hippocampus region analyzed. (g) Representative immunofluorescence images of GFAP (red fluorescence) and (h) quantitative analyses of GFAP mean fluorescence intensity in hippocampus. (i) Representative immunofluorescence images of Iba-1 (red fluorescence) and (j) quantitative analyses of Iba-1 mean fluorescence intensity in hippocampus. (k) Representative Western blots showing expression of GFAP, Iba-1, PSD-95, Synaptophysin and GAPDH. (l-q) Quantification of protein expression normalized to GAPDH. Data presented as mean ± SD. ****p<0.0001,***p<0.001,,**p<0.01, *p < 0.05, ns = not significant. Schematic created with BioRender.com).

**Fig. 11.**
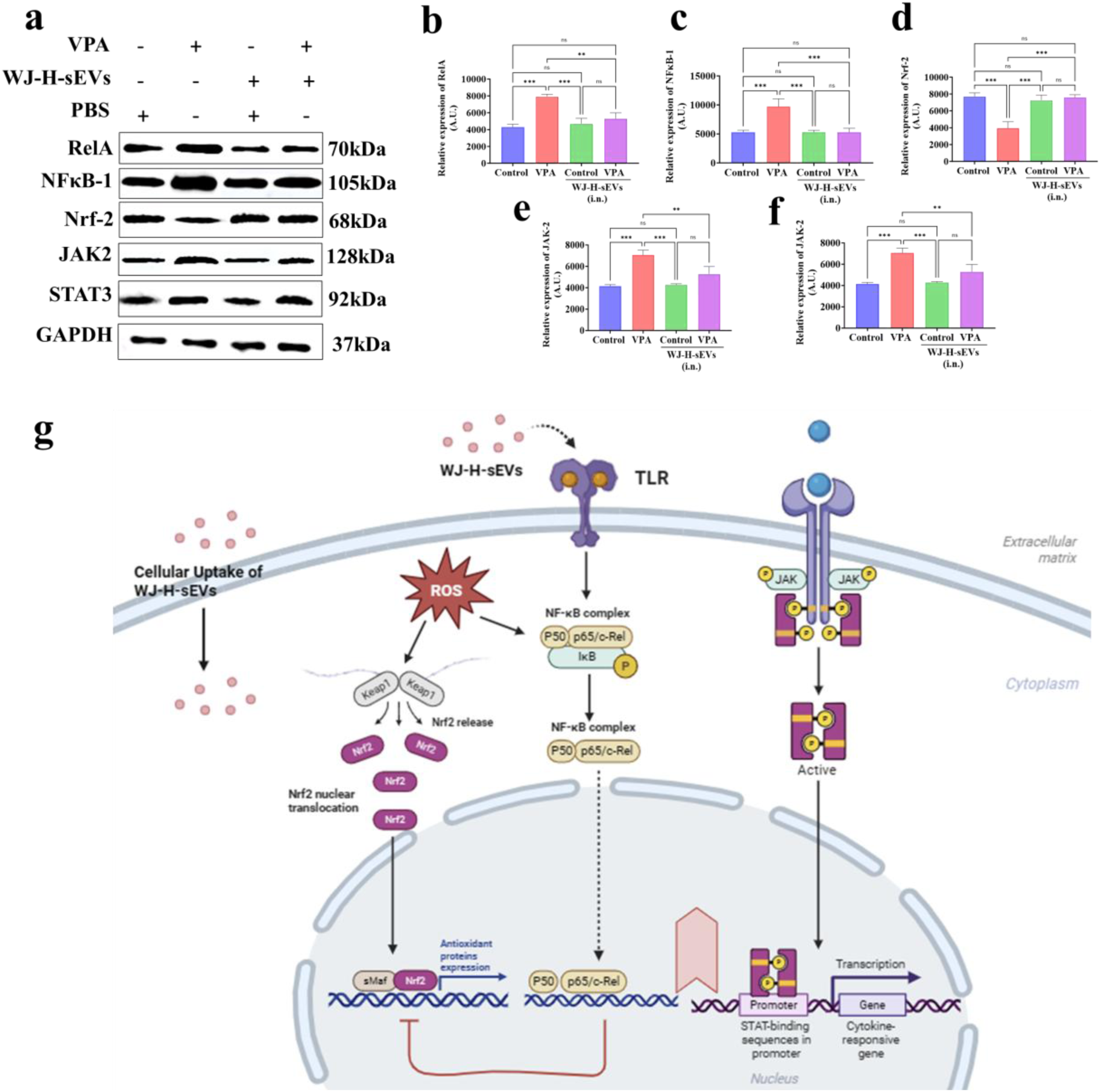
WJ-H-sEVs restore redox and neuroinflammatory balance in VPA-exposed mouse brains. (a) Representative Western blots showing expression of NF-κB subunits (RelA, NF-κB1), glial activation markers (Iba-1, GFAP), Nrf2, JAK2, and STAT3 in control, VPA, VPA + WJ-H-sEV, and control + WJ-H-sEV groups. (b-f) Quantification of protein expression normalized to GAPDH. (g) Schematic illustration of the proposed mechanism: WJ-H-sEV uptake reduces ROS-driven NF-κB and JAK2/STAT3 activation while enhancing Nrf2-dependent antioxidant signaling, collectively suppressing neuroinflammation and supporting synaptic integrity in VPA-induced ASD. Data presented as mean ± SD. ****p<0.0001,***p<0.001,,**p<0.01, *p < 0.05, ns = not significant. Schematic created with BioRender.com)

Given the central role of JAK/STAT signaling in sustaining neuroinflammation, we further broadened our analysis by assessing the expression of JAK1 and STAT3 expression. VPA-exposed mice exhibited marked upregulation of both JAK1 and STAT3, in agreement with previous preclinical reports implicating JAK/STAT signaling in the perpetuation of neuroinflammation and neuronal dysfunction in ASD models such as BTBR T+ Itpr3tf/J and VPA-induced ASD mice. Molecular intervention with WJ-H-sEVs significantly downregulated JAK1 and STAT3 expression, highlighting the vesicles capacity to exert broad regulatory effects on chronic inflammatory pathways. VPA also decreased Nrf2 expression and increased JAK2 and STAT3 (p < 0.01 vs control; Fig. 8a,d–f), whereas WJ-H-sEVs restored Nrf2 and significantly lowered JAK2 and STAT3 in VPA brains (p < 0.01 vs VPA; ns vs control + WJ-H-sEVs), consistent with rebalanced redox and cytokine-signalling axes (**Fig. 9a,d–f**).

A key observation of this study was the unique dual regulatory ability of WJ-H-sEVs to supress NF-κB/JAK-STAT and upregulate Nrf2 thereby establishing an antioxidant microenvironment capable of counteracting chronic neuroinflammation and oxidative stress, thereby shifting the neural milieu toward homeostatic balance.

The schematic in **Fig. 9g** summarizes a plausible mechanism whereby cellular uptake of WJ-H-sEVs dampens ROS-driven NF-κB and JAK2/STAT3 activation while enhancing Nrf2-dependent antioxidant gene expression, thereby reducing neuroinflammation and preserving synaptic architecture in VPA-induced ASD. Overall, these results demonstrate that WJ-H-sEVs uniquely achieve dual molecular regulation, positioning them as promising adjunctive therapeutics capable of targeting core pathogenic mechanisms inaccessible to current behavioral and pharmacological interventions. Given mounting evidence that neuroinflammation contributes to behavioral severity in ASD, such molecular modulation may offer meaningful improvements in long-term functional outcomes and quality of life.

## Discussion

As documented earlier by our group and other studies, given that MSCs naturally reside in low-oxygen microenvironments *in vivo*, hypoxic preconditioning may more accurately recapitulate their native niche, thereby preserving regenerative properties that are otherwise attenuated under standard normoxic culture conditions. A growing body of preclinical evidence across diverse animal models supports the therapeutic efficacy of hypoxia-conditioned MSCs and their secretome. In non-neurological models, hypoxia-primed MSCs or their secreted factors have shown superior outcomes in ischemic injury, inflammatory lung disease, myocardial infarction, and wound healing, largely attributed to enhanced angiogenic, immunomodulatory, and cytoprotective signaling. Similarly, in neurological and neurodegenerative models, including stroke, traumatic brain injury, spinal cord injury, and neuroinflammation, hypoxia-conditioned MSCs and their sEVs have been reported to improve neuronal survival, promote synaptic plasticity, modulate neuroinflammation, and enhance functional recovery compared to normoxia-conditioned counterparts. The clinical feasibility of hypoxia-conditioned MSC secretomes is further supported by a registered clinical trial in severe COVID-19 patients (NCT04374742), in which administration of hypoxia-primed MSC secretome demonstrated an acceptable safety profile and therapeutic benefit, providing translational proof-of-concept for this approach and highlighting its scalability and safety for manufacturing and therapeutic deployment of hypoxia-primed MSC-derived products in humans.

In our study, WJ-MSCs consistently outperformed BM-MSCs under both normoxic and hypoxic conditions, producing higher vesicle yields and exhibiting greater responsiveness to hypoxic conditioning. This likely reflects the intrinsic biological advantages of WJ-MSCs, including higher metabolic activity, shorter population doubling time, and a developmental origin that favors secretory and regenerative phenotypes. In line with the previous reports, the present results suggest that this strategy is not limited to a single MSC source and could be generalised to other tissue-derived MSCs to upscale vesicle yields for clinical applications.

Beyond quantity, hypoxia induced marked qualitative enhancements in sEV cargo, particularly in WJ-MSC-sEVs. Proteomic study,

The selective enrichment of miR-125b-5p, miR-145-5p, and miR-21-5p in WJ-H-sEVs than in BM-H-sEVs, points to a tissue-of-origin bias in miRNA packaging that is further amplified by hypoxia priming. These miRNAs are implicated in regulating neuronal survival, synaptic plasticity, glial activation and antioxidant defences, and are increasingly recognised as regulators of Nrf2-Keap1, JAK/STAT, and NF-κB pathways which are recognised as critical modulators of neuroimmune homeostasis. The preferential enrichment of this “neurorestorative” miRNA signature in WJ-H-sEVs provides a mechanistic basis for their superior functional performance and highlights the importance of both tissue origin and microenvironmental conditioning in EV-based therapies.

Consistent with this cargo advantage, WJ-H-sEVs displayed markedly superior uptake and pro-regenerative effects in C17.2 neural stem cells relative to BM- or normoxic sEVs. WJ-H-sEVs were internalised more efficiently, and robust pro-neuroregenerative effects and anti-inflammatory effects were observed *in vitro*. Synaptic transmission and plasticity abnormalities have been observed in the basal ganglia of the ASD mouse model. Given that ASD is characterised by altered neurogenesis, E/I imabalnce and widesperead synaptic and basal ganglia plasticity defects. These *in vitro* findings align with the miRNA profile and with broader literature showing that MSC-EVs can promote neural stem-cell proliferation, neurite outgrowth, and fate specification by delivering miR-21, miR-125 family members. The therapeutic effects of MSC derived sEVs in ASD models are increasingly linked to their miRNA cargo, which modulates synaptic plasticity, neuroinflammation, and neuronal survival.

After demonstrating that WJ-H-sEVs preferentially drive neuronal differentiation of C17.2 neural stem cells, we next asked whether these vesicles could also mitigate the oxidative and inflammatory stresses that are known to disrupt synaptic plasticity in ASD. Oxidative stress and neuroinflammation impair long-term potentiation, alter glutamatergic and GABAergic balance, and damage synaptic proteins transporters, thereby degrading circuit function in brain regions implicated in ASD, including hippocampus and basal ganglia. Many studies collectively show that patient-derived lymphocytes and fibroblasts from individuals with ASD exhibit reduced glutathione-dependent antioxidant defenses has been linked to impaired mitochondrial enzyme activity (e.g., decreased aconitase activity), excess mitochondrial superoxide production, neuroinflammation, immune dysregulation, and oxidative damage to macromolecules^96,97^. In this context, preserving mitochondrial function and redox balance is likely a prerequisite for sustained neuroregeneration and synaptic repair.

To probe this dimension, the antioxidant potential of hypoxia-primed sEVs was evaluated in both C17.2 neural stem cells and N9 microglia. WJ-H-sEVs more effectively preserved mitochondrial membrane potential and reduced mitochondrial superoxide accumulation than their normoxic counterparts and BM-derived sEVs, indicating superior protection against oxidative injury. In parallel, WJ-H-sEVs promoted a homoeostatic microglial phenotype, reducing pro-inflammatory polarisation and supporting a shift away from a chronically activated state. These findings support a dual mode of action in which WJ-H-sEVs not only foster neuroregenration but also stabilise the redox and inflammatory milieu in which those neurons must function.

Intranasal administration emerged as a clinically attractive delivery route, enabling direct brain access via olfactory and trigeminal pathways while minimizing systemic exposure. Compared with intravenous delivery, intranasal WJ-H-sEVs demonstrated preferential and sustained brain accumulation with reduced hepatic and splenic uptake. This enhanced CNS bioavailability likely reflects receptor-mediated uptake and membrane fusion mechanisms, further augmented by hypoxia-induced surface modifications such as increased LAMP-2b expression. Together, these features support intranasal delivery as a non-invasive, brain-targeted strategy suitable for repeated administration in pediatric and chronic neurodevelopmental disorders.

Using the prenatal valproic acid (VPA) model, which reliably recapitulates core behavioral and molecular features of ASD, intranasal WJ-H-sEVs produced robust rescue of anxiety-like behavior, exploratory deficits, recognition memory, and hippocampal-dependent spatial learning. Improvements across open field, novel object recognition, and Morris water maze paradigms reflected genuine cognitive recovery rather than nonspecific locomotor activation. These behavioral gains are consistent with restoration of synaptic plasticity within hippocampal–prefrontal circuits and compare favorably with prior MSC-EV studies in neurodegeneration, extending their relevance into ASD using a clinically tractable route.

The observed cognitive rescue is consistent with the known capacity of hypoxia-primed WJ-MSC-sEVs to mitigate oxidative stress and restore synaptic plasticity, both of which are profoundly disrupted in the VPA model. Oxidative stress is a central driver of synaptic dysfunction in ASD, where excessive reactive oxygen and nitrogen species impair long-term potentiation (LTP) by disrupting NMDA receptor redox sensitivity, calcium-dependent signaling cascades, and dendritic spine remodeling. By attenuating oxidative burden and neuroinflammatory signaling, WJ-H-sEVs may indirectly enhance glutamate clearance and normalize synaptic excitability, thereby supporting learning- and memory-related circuit function.

Collectively, these findings suggest that the behavioral recovery observed following WJ-H-sEV treatment arises from coordinated modulation of redox balance, synaptic plasticity, and excitatory–inhibitory homeostasis—mechanisms that are well established as convergent pathological drivers in ASD. The superior efficacy of hypoxia-primed WJ-sEVs further underscores the importance of tissue source and microenvironmental conditioning in optimizing sEV cargo for neurodevelopmental disorder intervention.

Oxidative stress and chronic inflammation (oxinflammation) are now widely recognized as central, interlinked drivers of ASD pathophysiology rather than secondary epiphenomena. Evidence from patient-derived immune cells, fibroblasts, post-mortem brain tissue, and peripheral biomarkers consistently demonstrates persistent redox imbalance coupled with sustained immune activation. Individuals with ASD exhibit impaired glutathione-dependent antioxidant capacity, mitochondrial dysfunction, excessive reactive oxygen species production, and increased oxidative damage to lipids, proteins, and nucleic acids. These abnormalities are accompanied by elevated pro-inflammatory cytokines, particularly IL-6, which has been shown to promote excitatory synapse formation while impairing inhibitory synapse development, thereby contributing to excitatory–inhibitory imbalance and synaptic dysfunction in ASD.

Consistent with this literature, our study demonstrates that WJ-H-sEVs exert coordinated anti-inflammatory and antioxidant effects in the VPA model. VPA exposure induced robust astrocytosis and microgliosis in the prefrontal cortex and hippocampus, elevated serum IL-6 with concomitant IL-10 reduction, and hyperactivation of NF-κB and JAK2/STAT3 signaling alongside suppression of Nrf2-dependent antioxidant pathways. Treatment with WJ-H-sEVs markedly attenuated GFAP⁺ astrocytosis and Iba1⁺ microgliosis, normalized IL-6/IL-10 ratios, suppressed NF-κB and JAK2/STAT3 activation, and uniquely restored Nrf2 signaling. These molecular changes were accompanied by preservation of synaptic proteins PSD-95 and synaptophysin, underscoring their functional relevance.

Mechanistically, NF-κB and JAK/STAT signaling act as convergent inflammatory hubs in ASD, sustaining chronic glial activation and cytokine production, while impaired Nrf2 signaling exacerbates oxidative stress and synaptic vulnerability. WJ-H-sEV treatment restored neuroimmune balance by simultaneously suppressing inflammatory cascades and re-establishing antioxidant defense. The superior efficacy of hypoxia-primed WJ-sEVs compared with normoxic counterparts suggests optimization of vesicular cargo, including miRNAs known to constrain NF-κB–dependent transcription and promote homeostatic glial phenotypes.

Together, these findings provide coherent molecular and histological evidence that WJ-H-sEVs dampen neuroinflammation, rebalance redox signaling, and preserve synaptic architecture through multi-node modulation of NF-κB, JAK2/STAT3, and Nrf2 pathways. This integrative mechanism is particularly well suited to the multifactorial biology of ASD, where convergent regulation of oxidative stress, immune activation, and synaptic stability is likely required to achieve meaningful behavioral recovery.

Direct comparison of BM- and WJ-derived sEVs under normoxic and hypoxic conditions demonstrates that MSC-sEV therapeutic efficacy is not uniform but is critically shaped by tissue origin and physiological priming. Our findings show that WJ-derived sEVs, particularly following hypoxic conditioning, are functionally superior to BM-derived counterparts, challenging the prevailing assumption that MSC-sEVs are interchangeable. Hypoxia not only increased sEV yield but also fundamentally reshaped vesicular cargo and biological potency, resulting in coordinated cellular, molecular, and behavioral rescue in an ASD model. These data underscore the importance of systematic source and conditioning optimization prior to clinical translation and position WJ-H-sEVs as a particularly potent and translationally attractive neurotherapeutic platform for ASD.

From a translational perspective, WJ-H-sEVs offer several advantages, including ready availability from discarded umbilical cords, minimal ethical concerns, favorable immunological properties, and scalability compatible with current regenerative medicine manufacturing pipelines. Our findings therefore provide strong empirical justification for prioritizing WJ-H-sEVs in early-phase ASD trials and support the integration of hypoxic priming into standardized sEV production workflows to maximize therapeutic efficacy.

Several limitations warrant consideration. This study employed a single environmental ASD model, and validation in complementary genetic models (e.g., *Shank3*, *CNTNAP2*) will be necessary to establish generalizability. In addition, while key signaling pathways—including NF-κB, JAK2/STAT3, and Nrf2—were modulated by WJ-H-sEVs, attribution to specific vesicular miRNAs or proteins remains inferential and will require targeted gain- and loss-of-function approaches. Finally, longer-term studies are needed to assess durability, dosing frequency, and safety of repeated intranasal administration.

Importantly, MSC-sEVs should be viewed not as replacements for existing behavioral or pharmacological therapies, but as biologically grounded adjunctive interventions capable of targeting oxinflammatory and synaptic mechanisms that remain largely unaddressed in ASD. By simultaneously dampening neuroinflammation, reducing oxidative stress, and stabilizing synaptic architecture, sEVs hold potential for disease-modifying effects. The intranasal route further enhances translational appeal by enabling non-invasive brain delivery with minimal systemic exposure, an approach already under clinical evaluation in pediatric populations.

Beyond ASD, the present findings have broader implications for the rational development of sEV-based therapeutics across neurodevelopmental and neurodegenerative disorders. While validated here in an ASD model, the source-agnostic benefits of hypoxic priming suggest that neuroregenerative efficacy can be enhanced even when WJ-MSCs are not available, providing flexibility for clinical translation. As a non-invasive, intranasally deliverable, and biologically targeted adjunct to existing behavioral interventions, MSC-sEVs offer a promising avenue to move beyond purely supportive care toward mechanism-informed, disease-modifying therapies. Future studies focusing on clinical validation, long-term safety, and cargo-specific mechanisms will be essential to accelerate the translation of this emerging therapeutic modality.

## Conclusion & Future Prospects

While prior investigations mainly exploited hypoxia to boost sEV yield or neuroregenerative potency, the present study establishes a definitive benchmark demonstrating that WJ-sEVs inherently possess richer cargo and superior functional competence than BM-sEVs under both normoxic and hypoxic conditions. Building on this advantage, hypoxic priming further amplifies the therapeutic potential, yielding WJ-H-sEVs that most effectively alleviate ASD-like behavioral, molecular, and neuroimmune abnormalities. Collectively, these findings position WJ-H-sEVs as a safe, non-invasive, cell-free adjunctive with the potential to be introduced in early neurodevelopmebtal years neurotherapeutics for ASD and support their advancement towards early-phase clinical evaluation. In addition, this work highlights discarded perinatal tissue as a sustainable clinical source of MSC-sEVs and establishes hypoxia as a scalable priming strategy that can be integrated into GMP-compatible manufacturing pipelines and extrapolated across other tissue-derived MSC sources to holistically refine and optimize sEV-based therapeutics.

