## Supplementary Tables for "Intranasal Delivery of Hypoxia-Primed Wharton’s Jelly MSC-Derived sEVs Reprograms Neuroimmune Signalling and Ameliorates Behavioural Deficits in a Valproic Acid–Induced Mouse Model of Autism Spectrum Disorder"

**Table S1. List of primary antibodies used in the study**

| **S.No.** | **Antibody** | **Company** | **Catalogue Number** |
| --- | --- | --- | --- |
| 1. | CD105 | eBiosciences, USA | 17-1057 |
| 2. | CD73 | Becton Dickinson, USA | 550257 |
| 3. | CD29 | eBiosciences, USA | 14-0299 |
| 4. | CD90 | Becton Dickinson, USA | 561558 |
| 5. | HLA-ABC | Becton Dickinson, USA | 555555 |
| 6. | HLA-DR | Becton Dickinson, USA | 555560 |
| 7. | CD34/45 | Becton Dickinson, USA | 341071 |
| 8. | CD63 | Abcam, USA | ab59479 |
| 9. | ALIX | Genetex, USA | GTX135282 |
| 10. | Calnexin | Genetex, USA | GTX13504 |
| 11. | GAPDH | Genetex, USA | GTX100118 |

**Supplementary Tables**

**Table S1. List of primary antibodies used in the study**

| **S.No.** | **Antibody** | **Company** | **Catalogue Number** |
| --- | --- | --- | --- |
| 1. | CD105 | eBiosciences, USA | 17-1057 |
| 2. | CD73 | Becton Dickinson, USA | 550257 |
| 3. | CD29 | eBiosciences, USA | 14-0299 |
| 4. | CD90 | Becton Dickinson, USA | 561558 |
| 5. | HLA-ABC | Becton Dickinson, USA | 555555 |
| 6. | HLA-DR | Becton Dickinson, USA | 555560 |
| 7. | CD34/45 | Becton Dickinson, USA | 341071 |
| 8. | CD63 | Abcam, USA | ab59479 |
| 9. | ALIX | Genetex, USA | GTX135282 |
| 10. | Calnexin | Genetex, USA | GTX13504 |
| 11. | GAPDH | Genetex, USA | GTX100118 |
| 12. | iNOS | eBiosciences, USA | 53-5920-82 |
| 13. | CD206 | eBiosciences, USA | 25-2069-42 |
| 14. | Arginase 1 | eBiosciences, USA | 17-3697-82 |
| 15. | CD14 | Abcam, UK | ab63319 |
| 16. | CD11b | eBiosciences, USA | 25-0118-42 |
| 17. | IL-6 | Affinity Biosciences, USA | DF6087 |
| 18. | IL-6R | Affinity Biosciences, USA | DF6466 |
| 19. | Rel-A p65 | Affinity Biosciences, USA | AF5006 |
| 20. | NFκB-1 p105/p50 | Affinity Biosciences, USA | AF6217 |
| 21. | TGFB2 | Affinity Biosciences, USA | AF0260 |
| 22. | SMAD4 | Affinity Biosciences, USA | AF5247 |

**Table S2 List of miRNA primers**

| **S.No.** | **miRNA primer** | **Company and Catalogue Number** | **GeneGlobe ID** |
| --- | --- | --- | --- |
| 1 | hsa-miR125b-5p | Qiagen, 339306 | YP00205713 |
| 2 | hsa-miR145a-5p | Qiagen, 339306 | YP00204483 |
| 3 | hsa-miR34a-5p | Qiagen, 339306 | YP00204486 |
| 4 | hsa-miR21-5p | Qiagen, 339306 | YP00204230 |
| 5 | U6 snRNA | Qiagen 339306 | YP02119464 |

**Table S1. List of primary antibodies used in the study**

| **S.No.** | **Protein** | **Company** | **Catalogue Number** | **Dilution used for Western Blotting Analysis** |
| --- | --- | --- | --- | --- |
| 1 | ALIX | GeneTex | GTX135282 | 1:500 |
| 2 | CD63 | ThermoFischer | 10628D | 1:1000 |
| 3 | Calnexin | GeneTex | GTX13504 | 1:3000 |
| 4 | GAPDH | GeneTex | GTX100118 | 1:3000 |
| **MSCs Characterization Antibodies** | | | | |
| 1 | CD105 | eBiosciences, USA | 17-1057 | 1:100 |
| 2 | CD73 | Becton Dickinson, USA | 550257 | 1:100 |
| 3 | CD29 | eBiosciences, USA | 14-0299 | 1:50 |
| 4 | CD90 | Becton Dickinson, USA | 561558 | 1:200 |
| 5 | HLA-ABC | Becton Dickinson, USA | 555555 | 1:100 |
| 6 | HLA-DR | Becton Dickinson, USA | 555560 | 1:100 |
| 7 | CD34/45 | Becton Dickinson, USA | 341071 | 1:100 |
